# Language Immersion Enhances Attentional Speech Processing via Low-Frequency Neural Tracking

**DOI:** 10.64898/2026.06.15.731669

**Authors:** Jingkang Wang, Taomei Guo, Mirjana Bozic

## Abstract

Language experience is a powerful driver of neurocognitive plasticity, shown to modulate attentional and executive processing in speakers of multiple languages. This paper investigated how immersion in a second-language environment shapes attentional processing of speech. Fifty-eight bilingual Chinese-English speakers matched on their English language proficiency listened to competing continuous speech streams in a naturalistic listening task. They were immersed either in their native language environment (Beijing, China), or in the second language environment (Cambridge, UK). In an identical EEG experiment across the two immersion contexts, we assessed the listeners’ cortical tracking of attended and unattended speech and task-related attentional allocation using Temporal Response Function (mTRF) and Power Spectral Density (PSD) analyses. Behavioral comprehension of the attended narratives was uniformly high. PSD analyses showed no group difference in task-related attentional allocation, but mTRF results revealed robust differences in cortical tracking, with increased tracking of attended - but not unattended - streams in the immersed group, driven by the delta band (1–4 Hz). This boost in tracking of the attended signal declined with prolonged immersion, indicative of changes to attentional speech processing as the language environment stabilizes and consistent with the expansion-renormalization framework of neurocognitive adaptation. Jointly, these data imply that immersion in second-language environments shapes the way listeners encode speech, sharpening the brain’s ability to extract target auditory information from background noise. They furthermore suggest that, rather than being static or monotonous, this modulation reflects a flexible and dynamic process that is continuously shaped by changes in environmental demands and their duration.

**Key points:**

1. Second language immersion boosts cortical tracking of attended speech, driven by the delta band (1–4 Hz).
2. This boost decreases with prolonged immersion, reflecting the dynamic, expansion-renormalization adaptation trajectory.
3. Immersion does not influence task-related attentional allocation.

## 1. Introduction

The human brain is a fundamentally adaptive system that continuously reorganizes its functional and structural architecture in response to environmental demands and experiential inputs (Zatorre et al., 2012; Sampaio-Baptista & Johansen-Berg, 2017). This neuroplasticity persists across the lifespan, with neuroimaging evidence showing experience-driven changes in both grey and white matter. For example, adults enrolled in a 9-month intensive second-language course showed progressive reorganization of white matter across multiple sites over the learning period (Schlegel et al., 2012), while right-handed participants learning left-hand writing showed training-related grey matter changes in primary motor cortices (Wenger et al., 2017a), providing direct evidence that neural structure can be recalibrated by experience. Comparable results are also observed in neural functions: training on a manual task over multiple days triggered widespread changes in frontoparietal activation patterns (Ruitenberg et al., 2018) while balance training led to reduced cortical activity in areas relevant for postural stability (Ruffieux et al., 2018). Crucially, this experience-based neuroplasticity is dynamic, with the magnitude and expression of neural modification depending on the type, intensity, and duration of experiences or environmental demands. One of the key models that formalizes this point is the expansion–renormalisation model of structural plasticity (Wenger et al., 2017b). It proposes that acquiring demanding skills, such as motor learning, tool-use learning and musical training, can induce dynamic changes in corresponding brain morphology, which demonstrates a nonlinear trajectory. Early stages of demanding learning can produce local expansion that reflects the exploration or formation of new neural pathways, followed by renormalisation as the amount of experience accumulates and expertise grows. The system eventually prunes or refines toward more efficient circuit configurations for the long-term skills (Quallo et al., 2009; Wenger & Kühn, 2020). Together, these findings illustrate that neurocognitive adaptations are shaped by the quantity and quality of experience, and by the sustained control demands that experience places on the system. In this paper, we apply the experience-driven framework to investigate the neurocognitive adaptations resulting from language immersion – a sustained exposure to a non-native linguistic environment – focusing on its effects on attentional processing of speech in bilingual listeners.

### 1.1 Language experience as a powerful driver of adaptation

Among the many forms of long-term experiences shaping the brain, bilingual language usage offers a unique perspective. The continuous engagement with multiple languages is a potent driver of neural plasticity, particularly within attentional and executive control networks in the frontoparietal areas (Tao et al., 2021). The altered attentional mechanisms in bilingualism stem from the joint activation of multiple languages (Marian et al., 2003; Martin et al., 2009; Starreveld et al., 2014), creating additional lexical competition and selection among cross-language alternatives compared to the lexical retrieval from a single set of candidates in monolinguals. The constant need to select the target language and inhibit the non-target language is therefore a plausible key driver of attentional and inhibitory adaptations in bilinguals, enabling them to process competing stimuli efficiently and flexibly despite the additional load (Bialystok et al., 2009; Kroll & Bialystok, 2013; Phelps & Bozic, 2025).

The reshaping of bilinguals’ executive-control mechanisms across the lifespan is particularly prominent in the domain of selective attention, the cognitive process of sustaining focus on a specific stimulus or information while simultaneously suppressing non-target inputs (Lavie, 2010). In the visual ERP studies, it has been argued that smaller N2 amplitudes for bilinguals in incongruent Stroop trials suggest more efficient selective attention to task-relevant information, and that shorter P3 latencies in incongruent Flanker trials suggest faster stimulus categorization (Kousaie & Phillips, 2017). Sullivan et al. (2014) reported that English speakers with six months of introductory Spanish course showed increased Go/No-go P3 amplitude, indicating that even early L2 learning can modulate attentional resource allocation during response inhibition. Morales et al. (2015) showed that bilinguals made fewer errors than monolinguals on conditions involving inhibition control in the AX Continuous Performance Task (AX-CPT), accompanied by larger N2 and P3 responses in the stimulus stage, arguably indicating more efficient reactive control and response inhibition. Comparable results also emerge from the domain of auditory selective attention. For example, when participants attended to stories in their native language paired with different types of interference, data showed stable neural tracking of the attended speech across interference types in bilinguals (Olguin et al., 2019), but the tracking varying as a function of the type of interference in monolinguals (Olguin et al., 2018). Using a similar paradigm in children, Phelps et al. (2022) found that monolingual and bilingual children differed in their neural tracking of attended speech, despite comparable behavioral performance. These results were interpreted as indicating greater attentional flexibility in bilinguals, highlighting the neuroplastic adaptation of attentional mechanisms to complex language environments.

Yet, although much of the existing evidence for bilingual influence on neurocognitive processes comes from studies that employed a binary comparison between bilingual and monolingual groups, it is important to recognize that bilingual experience should not be conceived as a coarse categorization, but a complex, dynamic, multidimensional spectrum encompassing various facets (DeLuca et al., 2019a; Yow & Li, 2015). This is especially important given the assumption that the type and amount of processing demands may drive the form and magnitude of experience-dependent neuroplasticity. Bilingual populations diverge widely in their language histories and variables such as L2 proficiency (e.g., Grundy & Timmer, 2017), age of L2 acquisition (AoA, e.g., Luk et al., 2011) and language immersion (e.g., DeLuca et al., 2019b), all of which can act as moderators of bilinguals’ attentional and executive control. Moreover, bilingualism is a long-term, linguistically and cognitively complex phenomenon and its impact on the processing system is not static, with e.g., the increase in bilingual experience eliciting expansion-renormalization trajectory of the volume of brain structures such as caudate nucleus and the nucleus accumbens (Korenar et al., 2023). Bilingualism-induced modulation of attentional control mechanisms therefore likely reflects structural and functional adaptation to the ongoing environmental demands of managing competing languages in specific contexts, rather than reflecting a single, fixed bilingual effect (Green & Abutalebi, 2013; DeLuca et al., 2019a, 2020a, 2024). Therefore, bilingualism should be conceptualized as an experience-based neuroplasticity framework, shifting the focus from broad binary comparisons to how specific aspects of language experiences fine-tune the neural mechanisms of bilinguals’ cognitive processing.

### 1.2 Effects of language immersion

Within this experience-based perspective, immersion in a second-language environment represents a particularly powerful yet underexplored dimension of bilingual experience influence on attentional speech processing. Language immersion typically refers to prolonged, natural use of the second language (L2) in an environment where L2 is the main language of communication, giving individuals sustained opportunities for real-world input and output in that language (Genesee, 1987; Johnson & Swain, 1997). Research has shown that immersive language-learning environments not only facilitate gains in language proficiency but also impact the brain’s structural and functional plasticity (Marín-Marín et al., 2022). Through sustained bilingual use, the brain exhibits adaptive changes at multiple levels, including increased grey matter volume, enhanced white-matter connectivity, and reorganization of neural networks involved in attentional control, executive function and speech processing (Pliatsikas, 2017). For example, L2 immersion schooling of French-speaking children into English classes over three years was shown to enhance alerting, auditory selective attention and mental flexibility, but not necessarily interference inhibition, when compared to non-immersion counterparts (Nicolay & Poncelet, 2013). Purić et al. (2017) tested early Serbian-speaking children after one year of English immersion. While no effect of L2 immersion on inhibition and shifting abilities were found, data showed improved working memory in children with high daily L2 exposure (5h per day), compared to the low-exposure immersion (1.5h per day) or the monolingual control group. This suggests that, beyond immersion status alone, the amount of immersion time may be critical for the emergence of any changes to executive control. Xie & Dong (2021) found that Chinese-English bilinguals who had been studying abroad (i.e., immersed in an English-speaking country) exhibited better mental set shifting than matched young adult peers without L2 immersion experience (i.e., studying at home in mainland China) as measured by the Wisconsin Card Sorting Test (WCST). They however showed no differences in the Flanker Task, which taps into interference inhibition and selective attention. DeLuca et al. (2019a) tested length of L2 immersion and extent of L2 use at home and in social settings against a battery of MRI measures, including grey-matter volume, subcortical structures, white-matter diffusivity and resting-state connectivity. They found that longer residence in the L2-speaking country and greater everyday L2 use in immersive settings were associated with systematic reshaping of subcortical structures in language and control networks (e.g., nucleus accumbens, thalamus, caudate nucleus and putamen). In a longitudinal MRI study investigating the effects of immersion, DeLuca et al. (2019b) scanned sequential L2 English bilinguals three years apart. While showing no change in their L2 proficiency, the continued three year immersion in L2 correlated with increases in grey-matter volume in the left cerebellum, increased white-matter diffusivity in the left frontal regions and reshaping of subcortical morphology in left caudate nucleus, amygdala and bilateral hippocampi. Data also suggested that longer L2 immersion and earlier AoA lead to longitudinal cerebellar increase. These data confirm the effects of immersion length on neurocognitive systems and provide direct evidence that prolonged naturalistic immersion in a L2-dominat environment dynamically drives ongoing neurological plasticity across the lifespan, even in immersed adult bilinguals who have already acquired L2 at a proficient level.

### 1.3 Current study

As reviewed above, the existing work on bilingual immersion has provided robust evidence for adaptations of neurocognitive structure and resting-state networks that scale with length and intensity of L2 immersion. Behavioral outcomes of immersion are however much more divergent across different executive tasks, with evidence for enhancement of alerting and working memory but no evidence for changes in interference inhibition, and the impact on selective attention remaining unclear. While this could be due to possible participant variation in dimensions other than immersion (e.g., L2 acquisition or proficiency), it is also likely to reflect the usage of static or artificial selective attention paradigms that do not tap into the online, task-relevant attentional processes. The current study therefore probes the effects of L2 immersion on selective attention during naturalistic listening, combining behavioral and neuroimaging measures to investigate how immersion experience and its length shape the dynamic functional mechanisms of attentional language processing.

To this end we investigated continuous speech processing in early Chinese-English bilinguals living in either Beijing, China (L1-dominant, non-immersed) or in Cambridge, UK (L2-dominant, immersed); matched on age, L2 AoA and L2 proficiency, and exposed to an identical naturalistic auditory selective attention task. Following Olguin et al. (2019) we used a dichotic listening paradigm where participants attended to a continuous narrative in one ear while simultaneously ignoring a competing stream in the other, thus simulating real-life scenarios of speech processing under interference. The type of interference was manipulated to create four conditions featuring distractors of varying intelligibility, ranging from purely acoustic to linguistic, and allowing us to control the processing load required to separate the attended from the unattended stream. Moreover, and in contrast to much of the existing immersion research focusing on L2 processing, we specifically assessed the processing of participants’ native language in this selective attention context. This was both because of the evidence that demands of L2-dominant environment might boost L2 activation while reducing L1 access (Linck et al., 2009), thus creating a possible confound, and because it allowed us to explicitly assess if any immersion effects on attentional processing might be generalized across both bilinguals’ languages. Therefore, to establish how L2 immersion might impact attentional speech processing of bilingual Chinese-English listeners, we assessed behavioral comprehension and cortical speech tracking in immersed and non-immersed participants attending to continuous Chinese narratives. Within the immersed group, we also evaluated how the length of L2 immersion modulates the neurocognitive mechanisms of attentional speech processing.

As standard in cortical speech-tracking studies, we focused our analyses on low-frequency EEG signal (1-12 Hz) since this range captures slow cortical oscillations that most consistently entrain to the temporal envelope of continuous speech, either alone or in multi-talker, selective-attention environments (e.g., Attaheri et al., 2022; Ding & Simon, 2012; Etard & Reichenbach, 2019; Giraud & Poeppel, 2012; Obleser & Kayser, 2019; Zuk et al., 2021), while excluding higher-frequency EEG activity which is less directly tied to speech processing and more vulnerable to muscle movement and line noise (Muthukumaraswamy, 2013; Whitham et al., 2007, Leske & Dalal, 2019). Importantly, low-frequency neural entrainment is not functionally homogeneous within this range: delta-band (∼1-4 Hz) speech tracking is associated with top-down speech processing such as syntactic and semantic segmentation and integration, or effortful listening in challenging conditions; whereas theta-band (∼4-8 Hz) entrainment more closely reflects bottom-up acoustic, phonemic and syllabic structure of the speech signal (Bröhl & Kayser, 2021; Doelling et al., 2014; McHaney et al., 2021). Alpha band (∼8-12 Hz) activity, in contrast, is not typically tied to robust envelope tracking, but to speech intelligibility and noise suppression (Dimitrijevic et al., 2017; Foxe & Snyder, 2011; Obleser & Weisz, 2012; Strauß et al., 2014). Thus, any immersion group differences observed in broadband speech tracking (1-12Hz) could arise from modulations to different types of neurocognitive mechanisms, and identifying and separating these processes is critical for establishing how language immersion might affect attentional speech tracking in bilinguals. We thus also decomposed EEG signal into the canonical delta, theta and alpha frequency bands, and assessed frequency-specific effects to pinpoint which mechanisms of stimuli tracking (e.g., top-down or bottom-up processing) drive the potential modulations by L2 immersion.

Additionally, we also conducted spectral power analyses in these same bands, as power variations of neural oscillations index the strength of task-related neural engagement and resource allocation, where different bands are again linked to partially distinct cognitive functions and have well-established roles in auditory processing, especially for speech comprehension requiring attention (e.g., Buzsáki & Draguhn, 2004; Jensen & Mazaheri, 2010; Klimesch et al., 2007; Pfurtscheller & Lopes da Silva, 1999; Wöstmann et al., 2016). For instance, delta power is associated with slow, large-scale integrative processes, internal concentration or sustained attention to relevant inputs (Knyazev, 2007; Harmony, 2013). Theta power indexes working memory and cognitive control, increasing with conflict monitoring and maintenance demands (Cavanagh & Frank, 2014; Knyazev, 2007; Sauseng et al., 2010). Alpha-band power is a key marker of speech intelligibility, selective inhibition control and regulation of listening effort in difficult conditions (Foxe & Snyder, 2011; Frey et al., 2015; Obleser & Weisz, 2012; Wöstmann et al., 2016; 2021). Therefore, examining delta, theta and alpha EEG power alongside cortical speech tracking allowed us not only to measure how robustly the brain follows the speech envelope, but also to demonstrate how strongly large-scale networks for integration, control and selective inhibition are engaged by continuous L1 speech comprehension under sustained interference. Comparing power changes across bands between groups thus provides complementary insights into whether two immersion groups dedicate equal attention resources to the dichotic listening task.

## 2. Methods

### 2.1 Participants

A total of 62 participants took part in the experiment, with 31 bilingual participants recruited in Beijing, China (without English immersion) and 31 bilinguals recruited in Cambridge, UK (with English immersion) respectively. All participants were native Mandarin Chinese speakers aged 18-38 (mean age 23.1; 18 male) who acquired L2 English by the age of six, classifying them as early bilinguals (Berens et al., 2013). Participants first completed an online survey with demographic questions and adapted Bilingual Language Profile (BLP; Birdsong et al., 2012), which assesses bilinguals’ language history, use, proficiency, and attitude. They then completed LexTALE (Lemhöfer & Broersma, 2012), a 60-item lexical decision task that has been validated as an objective measure of general English proficiency. They were included in the experiment only if they reported learning English by the age of six, were fully proficient in both Mandarin and English, and scored ≥60% on the LexTALE test (equivalent to B2 or higher proficiency). All participants were healthy adults reporting no neurological or hearing problems.

All 62 participants completed the in-person EEG experiment and were reimbursed for their participation. Two participants were removed for channel noise, one was disqualified for attentional failure, indicated by below-chance performance on comprehension questions, and one was excluded for being more than 3 SD older than the mean, resulting in a final sample of 58 participants in total for two sites. Table 1 summarizes demographic and language background characteristics of participants across two testing sites, showing that participants at both sites were comparable in sex, age [*t*(56) = -0.789, *p* = .433], age of L2 acquisition (*W* = 361, *p* = .355) and L2 proficiency (*W* = 320.5, *p* = .128). BLP questionnaire responses for Cambridge and Beijing participants are given in the Supplementary Materials, Table S1.

**Table 1.** Participant information across testing sites (SD in brackets)

| Site | N | Female /Male | Age in years | Age range | AoA English (years) | LexTALE % |
| --- | --- | --- | --- | --- | --- | --- |
| Beijing | 31 | 23/8 | 22.39 (2.65) | 18-30 | 4.52 (1.36) | 76.90 (13.57) |
| Cambridge | 27 | 20/7 | 22.96 (2.90) | 18-29 | 4.89 (1.05) | 81.19 (10.10) |

### 2.2. Materials

Identical stimuli and procedures were applied to participants in both Cambridge and Beijing. The experiment employed a dichotic listening paradigm in which participants were instructed to attend to a continuous, naturalistic narrative presented in one ear while simultaneously ignoring a competing stream in the other. The design simulated real-life listening situations that demand selective attention to the target speech amidst distracting auditory inputs. Participants always attended to stories in their native language (Chinese), whereas the type of the interfering stream was systematically manipulated to create four experimental conditions featuring distractors of varying intelligibility: 1) **Single-Talker**, where participants were only presented with the attended Chinese narrative in one ear with no competing input; 2) **Native-Musical Rain (Chinese-MuR)**, where the distractor was a nonlinguistic signal that mimics the acoustic properties of prolonged speech without evoking linguistic perception 3) **Native-Unknown (Chinese-Serbian)**, where the interfering narrative was in Serbian, a language that listeners did not understand; and 4) **Native-Native (Chinese-Chinese)**, in which the distractor was a different Chinese narrative, producing the strongest interference. The participants’ L2 English was not part of any of the experimental conditions. A summary of the experimental conditions is displayed in Table 2.

**Table 2.** Summary of Experimental Conditions.

| Condition | Attended | Unattended |
| --- | --- | --- |
| Single Talker | Chinese (Native language) | No Interference |
| Native - MuR | Chinese (Native language) | Musical Rain (Non-linguistic) |
| Native – Unknown | Chinese (Native language) | Serbian (Unknown language) |
| Native – Native | Chinese (Native language) | Chinese (Native language) |

Stimuli included 10 Chinese children’s stories, two Serbian stories, and two Musical Rain (MuR) streams, with each transcribed into 120 sentences (∼3 seconds each). Chinese and Serbian stories were recorded by female native speakers respectively, with different Chinese speakers for attended and unattended stories. During testing, each story was presented in two consecutive blocks (first 60 vs. last 60 sentences), with 300-ms silent gaps between sentences. Musical Rain (MuR) stimuli were created by extracting temporal envelopes from Chinese narratives and filling them with randomized fragments of synthesized speech. This preserved the spectrotemporal properties of natural speech while eliminating continuous formants thus preventing the triggering of speech percept (Bozic et al., 2010; Uppenkamp et al., 2006). All stimuli were RMS-normalized to control sound amplitude.

### 2.3 Procedure

Before the experiment, all participants provided written informed consent, and the study was approved by the Institutional Review Board from both sites. Stimuli were delivered via MATLAB with Psychophysics Toolbox (Brainard, 1997; Pelli, 1997) through sound-insulated earphones in a quiet room. Each condition comprised 4 blocks (2 attended stories, 240 sentences), each lasting ∼3.2 minutes. In total, each participant heard 960 attended sentences and 720 interfering trials. The ‘Single Talker’ condition was always presented first for familiarization, while the remaining conditions and story order were randomized across participants, with attended narratives alternating between ears from block to block. The attended side in the first block (i.e., beginning with listening to the left or right side) was also randomized across participants.

To ensure participants attended to narratives as instructed, each block was followed by 10 true/false comprehension questions (e.g., ‘*The king ordered the servant to retrieve the ring from the sea’,* Y/N). The number of correct and incorrect statements was equal for each story. In total, each participant answered 160 comprehension questions in Mandarin Chinese on Qualtrics. During this self-paced section, participants were allowed to take a short break between blocks if needed. The experimental procedure is demonstrated in Figure 1a. Transcripts of all narratives and the full set of comprehension questions are provided in the OSF directory https://osf.io/gsrje/

**Figure 1.**
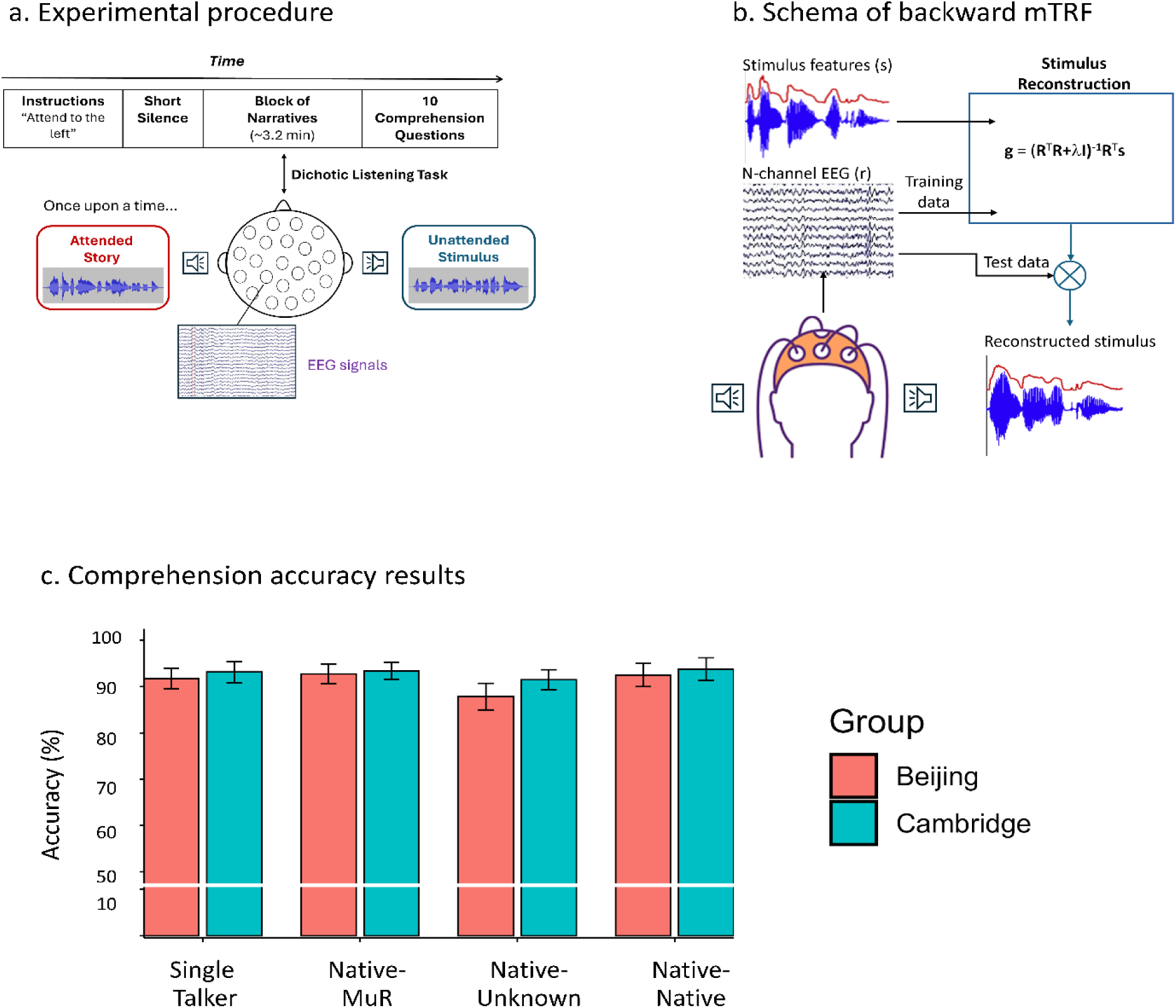
(a) Sequence of a block: Participants were instructed to attend to one side. Following short silence, narratives were presented for 3.2 min and participants were then asked to complete 10 comprehension questions about the attended story. (b) mTRF stimulus reconstruction: A backward mTRF decoding model was fit using a leave one-out cross-validation procedure. This generated a reconstruction of speech envelopes that were validated against the original envelopes. (c) Comprehension accuracy scores: Results showed that all participants performed the task well, with uniformly high scores across groups and conditions.

### 2.4 Electroencephalography (EEG) Data: Acquisition and Pre-processing

EEG data at both sites were acquired using the same Netstation Acquisition system with a high-density 128-channel Ag/AgCl electrode net (Electrical Geodesics, Inc.), and with Cz as the reference electrode. The sampling rate was 500 Hz. All raw EEG data were imported and pre-processed in MATLAB using the EEG toolbox EEGLAB (Delorme & Makeig, 2004). For each participant, time markers indicating the onset of each attended and unattended sentence were inserted into the EEG data to pinpoint the latency of its presentation. Each marker was labeled with the sentence status (attended/unattended), story index, experimental condition and the corresponding sentence number within the block. Data were band-pass filtered (1-40 Hz) to eliminate low-frequency drifts and high-frequency artifacts as well as line noise (Leske & Dalal, 2019), then resampled to 100 Hz for alignment with the speech envelopes of auditory signals. Bad channels were identified using power spectrum (-/+ 3 SDs) and kurtosis (-/+5 SDs), then rejected and interpolated using data from neighboring electrodes. Next, the EEG was re-referenced to the average of all channels (de Cheveigne & Arzounian, 2018). Afterwards, Infomax Independent Component Analysis (ICA) was applied to isolate underlying sources from the signal by maximizing their statistical independence (Subasi & Gursoy, 2010), and non-brain components such as eye blinks, muscle activity, and channel noise were removed. Based on the time markers, data were then segmented into 2.7s epochs (2.5 s sentence duration + 0.2 s pre-stimulus baseline) at the sentence level, yielding 960 attended and 720 unattended epochs per participant. Noisy epochs with poor signal quality were excluded.

### 2.5. Data Analysis

#### 2.5.1 Behavioral Data Analysis

To test whether the probability of correctly answering comprehension questions differed between the two immersion groups and across listening conditions, we ran a generalized linear mixed-effects model using the glmer function in R (Bates et al., 2015). The dependent variable was comprehension accuracy for each question, coded dichotomously (1 = correct, 0 = incorrect). Fixed effects were Immersion Group (two levels: Beijing and Cambridge participants), Condition (four levels: Single Talker, Chinese-MuR, Chinese-Sebian Chinese-Chinese), Group by Condition interaction, participant age, sex, education background and L2 English proficiency (LexTALE score). Participants and comprehension questions were included as random intercepts.

#### 2.5.2 EEG Data Analysis

The primary EEG analysis was the cortical tracking of speech, analysed using the Multivariate Temporal Response Function (mTRF) Toolbox in MATLAB (Crosse et al., 2016). The mTRF framework models the linear mapping between auditory stimuli (e.g., speech envelope) and neural responses (EEG) across time lags, thereby quantifying how the brain dynamically tracks the speech signals. For each attended and unattended stream, the speech envelope was extracted with the *Hilbert2* function (Ktonas & Papp, 1980), which captures rhythmic and prosodic cues by reflecting temporal fluctuations in signal energy (Muda et al., 2010; Tiwari, 2010). Envelopes were downsampled to 100 Hz to match the EEG data and zero-padded as necessary to align with EEG epoch lengths. The EEG data were filtered between 1-12 Hz, a frequency range that captures the low-frequency neural activity (delta, theta, alpha) most consistently associated with cortical tracking of continuous speech and auditory selective attention (e.g., Ding & Simon, 2012; Giraud & Poeppel, 2012; Obleser & Kayser, 2019), while filtering out less relevant higher-frequency noise. We adopted the backward (decoding) model, which reconstructs the auditory speech envelope from multichannel EEG responses: g = (R^T^R+λI)^-1^R^T^s where g is the decoder (i.e., mapping weights from EEG responses to stimulus), R is the EEG data matrix across channels, s is the envelope, λ is the ridge parameter, and I is the identity matrix (Crosse et al., 2016). The schematic of the backward mTRF model is shown in Figure 1b.

To assess how immersion in L2-dominant environment modulated the neural mechanisms of speech tracking under interference, we quantified the Pearson’s correlation (r) between the reconstructed speech envelopes decoded from the EEG signal and the original speech envelopes in each participant group. Correlations were averaged across trials and channels to yield a single score per participant for each listening condition. To generate the baseline distribution that represents chance-level reconstruction accuracy, we shifted the stimulus signal by a random amount (minimum ±2.5 s to avoid overlapping with the true lag window) to break alignment, and then computed the correlation (control r value) between the predicted and shifted envelope (Crosse et al., 2016, 2021). The paired sample *t*-test showed that the correlations between the real attended stimuli and signals reconstructed from EEG data were significantly higher than the chance-level for participants in both Cambridge [*t*(107) = 23.208, *p* < .001] and in Beijing [*t*(123) = 18.693, *p* < .001]), and this also held for the unattended streams [Cambridge: *t*(80) = 7.579, *p* < .001; Beijing: *t*(92) = 6.580, *p* < .001]. The correlations for real stimulus-to-EEG alignment were thus constantly higher than the control values, indicating that the decoding performance was above chance-level and that EEG captured the genuine time-locked information about the auditory signals in the model. In addition, to quantify the separation between real and control distributions, D-prime (*d*^′^) was calculated as a measure of sensitivity using the formula: 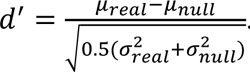. Across participants, the real decoding performance of attended stimuli was reliably separated from the control distribution in both Beijing (mean d′ = 2.075, SD = 1.062) and Cambridge (mean d′ = 3.176, SD = 1.070).

The attended and unattended correlation values were analyzed in R using linear mixed-effects models to test the effects of Attention, Condition, Immersion Group and their interactions. Participant was added as the random intercept. To delve into frequency-specific modulations, the same set of analysis was replicated for the decomposed EEG data in delta (1-4Hz), theta (4-8Hz) and alpha (8-12Hz) bands respectively. Finally, for Cambridge participants only, the length of time they have been living in an English-speaking country (L2 immersion) was added as a predictor in the linear mixed-effects model.

Spectral power analyses were conducted to investigate frequency-specific oscillatory EEG activity in delta, theta, and alpha ranges during the dichotic-listening task. The spectral power analyses were performed with FieldTrip (Oostenveld et al., 2011) using the multitaper time-frequency convolution method. Analyses focused on attended trials and the four listening conditions, in both Beijing and Cambridge groups. The pre-listening period in each block served as the baseline. EEG epochs were segmented into baseline and listening and matched in length (2.7 s). For each window, we computed spectra from 1-12 Hz in 1-Hz steps using 1-s Hanning tapers with a sliding step of 0.1 s. For each condition, power was averaged across time and channels to yield participant-level spectra for baseline and listening periods respectively. Baseline was subtracted from listening to obtain normalized spectra per condition that enables across-subject comparisons. We calculated band-limited power from the normalized spectra by averaging within canonical bands: delta (1-4 Hz), theta (4-8 Hz) and alpha (8-12 Hz). A linear mixed-effect model was used to test whether the normalized power in each frequency band is affected by listening conditions and immersion group.

## 3. Results

### 3.1 Behavioral measures

The mean response accuracy to comprehension questions was 92.96% (SD = 5.53%) in Cambridge and 91.21% (SD =6.90%) in Beijing, suggesting that all listeners focused on the target speech as instructed. The model testing whether the probability of correct answers differed across groups and conditions showed no difference across groups [*χ*^2^(1) = 0.923, *p* = .337] or conditions [*χ*^2^(3) = 5.373, *p* = .146], nor interaction between group and condition [*χ*^2^(3) = 2.768, *p* = .429]. The average comprehension accuracy by group and condition is illustrated in Figure 1c.

### 3.2 EEG: Cortical Speech Tracking

#### 3.2.1 Broadband tracking (1-12Hz)

The attended and unattended correlation *r* values obtained from the backward mTRF model are illustrated in Figure 2a. A linear mixed-effects model was fit to predict the *r* values from factors of Attention (attended, unattended), Immersion Group (Beijing, Cambridge), Interference Condition (Native-MuR, Native-Unknown, Native-Native), and their interactions. The random intercept for participants was included. The results revealed significant main effects of Attention [*F*(1, 280) = 430.603, *p* < .001, 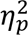 = .606] and Immersion Group [*F*(1, 56) = 12.532, *p* < .001, 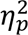 = .183] but not Condition [*F*(2, 280) = 2.594, *p* = .077, 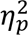 = .018]. Significant interaction was found between Attention and Group [*F*(1, 280) = 29.472, *p* < .001, 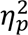 = .095] as well as between Attention and Condition [*F*(2, 280) = 31.204, *p* < .001, 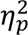 = .182]. There was no interaction between Condition and Group [*F*(2, 280) = 0.807, *p* = .447, 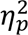 = .006]. Tukey-adjusted post-hoc comparisons showed that the tracking of attended stimuli was consistently stronger than that of unattended stimuli in both groups and across all three interference conditions, suggesting that the attended stimuli were generally tracked more robustly regardless of L2 immersion or the type of interference. Post-hoc tests also showed more robust tracking of attended stimuli in the Cambridge than the Beijing group in all three interference conditions (Native-MuR : Δ*M* = 0.035, *SE* = 0.006, *t*(223.61) = 5.466, *p* < .001; Native-Unknown: Δ*M* = 0.024, *SE* = 0.006, *t*(223.61) = 3.756, *p* < .001; Native-Native: Δ*M* = 0.021, *SE* = 0.006, *t*(223.61) = 3.244, *p* = .001; where ΔM signifies the mean decoding difference between Cambridge and Beijing groups). The tracking of unattended stimuli did not differ between groups in any condition (all *p* > .05), showing that the differences between groups were driven by the tracking of attended narratives only. The full lmer model results and all post-hoc comparisons are summarized in Table S2 in the Supplementary Materials.

**Figure 2.**
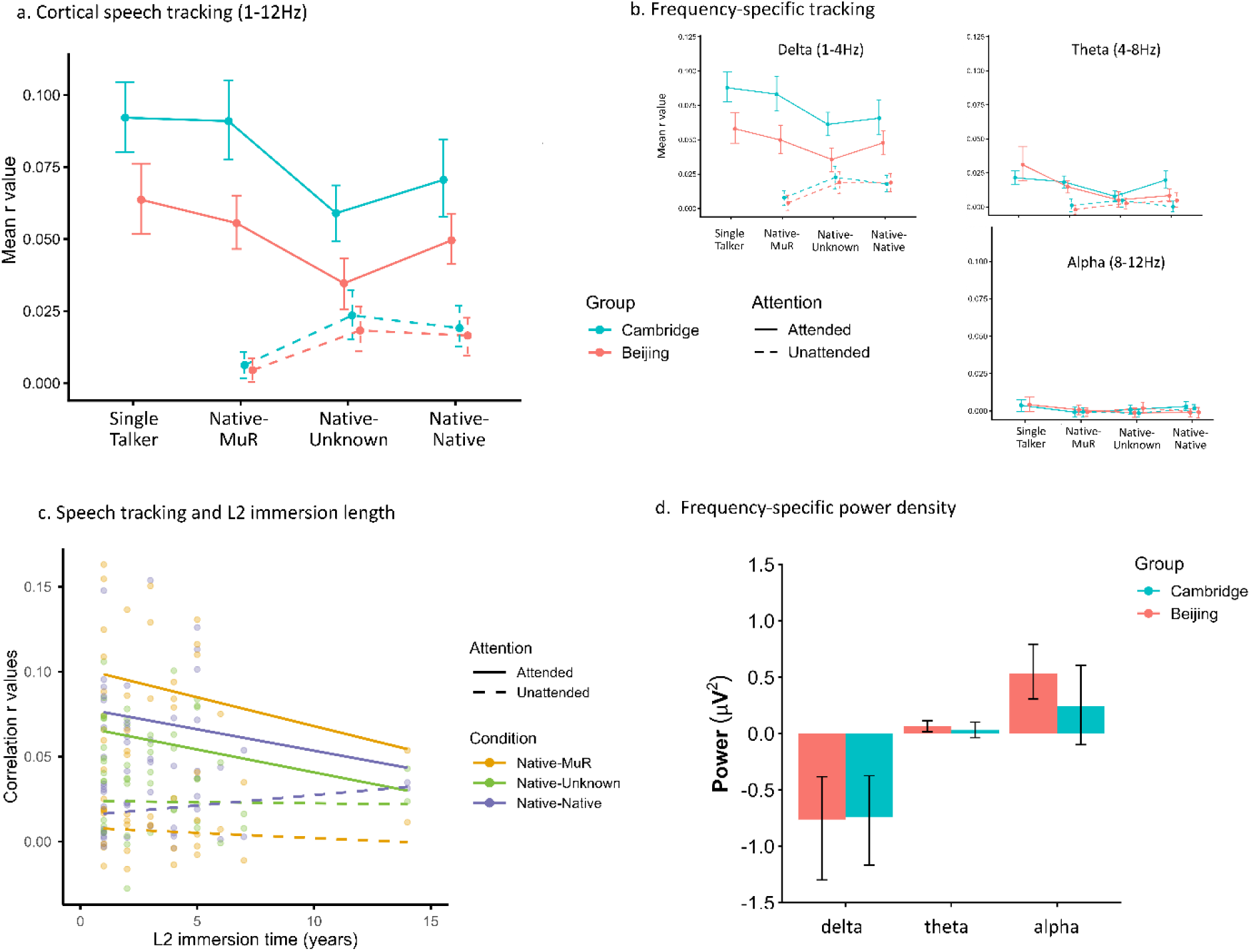
a) Broadband speech tracking results, revealing highly consistent patterns across the two groups, but with significantly more robust tracking of attended speech in the Immersed (Cambridge) group. (b) Frequency-specific tracking, illustrating that the effects were driven by the delta frequency range (1-4Hz). (c) Effects of L2 Immersion length on speech tracking in the Immersed group. (d) Power Spectral Density Analysis, showing comparable spectral power distribution across immersion groups in different frequency bands.

A follow up analysis focused on attended streams only, including Group (Beijing vs. Cambridge), all four Conditions (Single Talker, Native-MuR, Native-Unknown and Native-Native), their interaction and participants as the random intercept (Table S3, Supplement). It showed a significant main effect of Group [*F*(1, 56) = 17.446, *p* < .001, 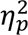 = .238] and Condition [*F*(3, 168) = 26.470, *p* < .001, 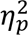 = .321], but no interaction [*F*(3, 168) = 1.298, *p* = .277, 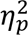 = .023]. The Tukey-adjusted post-hoc comparisons indicated that, while the attended steams paired with weaker interference were encoded more strongly than those paired with stronger interference in both groups, the attended encoding was consistently higher in Cambridge than in Beijing group in all conditions (Single Talker: Δ*M* = 0.029, *SE* = 0.008, *t*(119) = 3.534, *p* < .001; Native-MuR: Δ*M* = 0.035, *SE* = 0.008, *t*(119) = 4.392, *p* < .001; Native-Unknown: Δ*M* = 0.024, *SE* = 0.008, *t*(119) = 3.018, *p* = .003; Native-Native: Δ*M* = 0.021, *SE* = 0.008, *t*(119) = 2.607, *p* = .010).

The same linear mixed-effects model with predictors of Condition (3 levels) and Group was also fit for unattended streams. The result only showed a significant main effect of Condition [*F*(2, 112) = 13.574, *p* < .001, 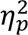 = .195], but no effects of Group [*F*(1, 56) = 0.989, *p* = .324, 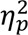 = .017] and no interaction [*F*(2, 112) = 0.159, *p* = .853, 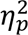 = .003]. Tukey-adjusted post-hoc tests revealed identical pattens in the two groups, such that the unattended MuR stimuli were tracked more weakly than unattended narratives in native and unknown languages for both Cambridge and Beijing groups [Native: Cambridge group Δ*M* = -0.013, *SE* = 0.005, *t*(112) = -2.785, *p* = .017; Beijing group Δ*M* = -0.012, *SE* = 0.004, *t*(112) = -2.793, *p* = .017; Unknown: Cambridge group Δ*M* = -0.017, *SE* = 0.005, *t*(112) = -3.736, *p* < .001; Beijing group Δ*M* = -0.014, *SE* = 0.004, *t*(112) = -3.210, *p* = .005]. The lmer results for unattended streams and the corresponding post-hoc tests are summarized in Table S4 in Supplementary Materials.

#### 3.2.2 Frequency Decomposition: Speech Tracking in Delta, Theta and Alpha ranges

The same mTRF analyses were replicated on delta (1-4 Hz), theta (4-8 Hz) and alpha frequency bands (8-12 Hz) separately. The attended and unattended *r* values by Condition and Immersion Group for each frequency band are displayed in Figure 2b. In the delta band (1-4Hz), the results fully replicated those observed in the broadband (1-12 Hz) analysis, with a significant main effect of Attention [*F*(1, 280) = 347.587, *p* < .001, 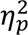 = .554] and Group [*F*(1, 56) = 11.221, *p* = .001, 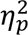 = .167], and interaction between Attention and Group [*F*(1, 280) = 26.981, *p* < .001, 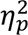 = .088] and Attention and Condition [*F*(2, 280) = 18.351, *p* < .001, 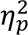 = .116]. No interaction between Group and Condition was found [*F*(2, 280) = 1.636, *p* = .197, 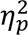 = .012]. Same held for the analysis of attended streams only, which demonstrated a significant main effect of Condition [*F*(3, 168) = 17.401, *p* < .001, 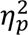 = .237] and Group [*F*(1, 56) = 16.147, *p* < .001, 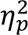 = .224], but no interaction [*F*(3, 168) = 1.622, *p* = .186, 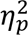 = .028]). Finally, the analysis of the unattended streams in the delta band indicated a significant main effect of Condition [*F*(2, 112) = 12.150, *p* < .001, 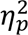 = .178] but not of Group [*F*(1, 56) = 0.405, *p* = .527, 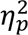 = .007] nor their interaction [*F*(2, 112) = .349, *p* = .707, 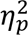 = 006]; again fully replicating the results from the 1-12 Hz band. In the theta band (4-8 Hz), the correlation values were generally much lower. Using the same analyses, data showed a significant main effect of Attention [*F*(1, 280) = 55.150, *p* < .001, 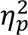 = .165], and significant interactions between Attention and Group [*F*(1, 280) = 4.149, *p* = .043, 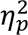 = .015] and Attention and Condition [*F*(2, 280) = 8.824, *p* < .001, 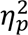 = .059]. The analysis of attended streams alone showed a main effect of Condition [*F*(3, 168) = 11.873, *p* < .001, 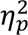 = .175] and interaction between Group and Condition [*F*(3, 168) = 3.320, *p* = .021, 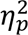 = .056], while the analysis of unattended channel showed no main effects or interactions (all *p* > .05). In the alpha band (8-12 Hz), the neural tracking of auditory stimuli was weak and did not significantly differ from the control distribution. Taken together, the neural tracking results from different frequency bands show that the decoding differences between groups in the dichotic listening task were primary driven by the delta band (1-4 Hz). The statistics of the lmer tests for delta and theta bands are summarized in Tables S5-10 respectively in the Supplement.

### 3.3 Effects of L2 Immersion Length on Speech Tracking

For the group of bilingual participants residing in Cambridge (N = 27), the length of time they have been immersed in an English (L2)-dominant language environment was extracted and added to a model including Attention (attended, unattended), Condition (Native-MuR, Native-Unknown, Native-Native) and their interaction. The L2 immersion time for bilinguals in Cambridge ranged from 1 to 14 years (Mean = 3.22, SD = 2.81). Results showed the main effect of Attention [*F*(1, 125) = 171.166, p < .001, 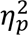 = .578] and the interaction between Attention and Condition [*F*(2, 125) = 8.047, p < .001, 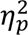 = .114], such that attended decoding was higher than unattended decoding, but this difference was moderated by Condition. While the main effect of L2 immersion time was not significant ([*F*(1, 25) =1.077, p = .309, 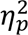 = .041], there was a significant interaction between L2 immersion time and Attention [*F*(1, 125) = 6.249, *p* = .014, 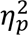 = .048], such that longer L2 immersion time led to weaker tracking of the attended stream. Figure 2c illustrates the fit regression lines depicting the relationship between r values and L2 immersion length in each condition, with the model output summarized in Table S11 in Supplementary Materials. Since one participant’s immersion time was notably longer than that for the rest of the group, we also replicated the same analysis excluding this participant. The results (Table S12 and Figure S1 in the Supplement) clearly show a comparable trend to the full dataset, although without a significant attention by immersion time interaction, which we return to in the Discussion.

### 3.4 Power Spectral Density Analysis

The normalized power spectral density (PSD) for each subject group is illustrated in Figure 2d. A model predicting the normalized PSD in 1-12 Hz from Immersion Group, Condition and their interactions did not show an effect of Group [*F*(1, 56) = .392, *p* = .534, 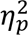 = .007] or Condition [*F*(3, 632) = .861, *p* = .461, 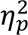 = .004], nor their interaction [*F*(3, 632) = 1.610, *p* = .186, 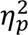 = .008]. To match the cortical speech tracking analyses performed above, we also performed the same analyses for delta, theta and alpha bands separately. Across all bands, there were no effects of group or condition, and no interactions. All PSD results are shown in Table S13 in Supplementary Materials. Taken together, these results suggest that the spectral power distribution was comparable across immersion groups, implying no difference in task-related attention allocation and neural engagement.

## 4. Discussion

This study investigated how immersion in an L2 context impacts bilinguals’ attentional processing of speech. Using the identical experimental setup, the study compared attentional speech processing in adult Chinese-English bilinguals residing in Cambridge as the immersed group and in Beijing as the non-immersed group, respectively. In a dichotic listening task, participants attended to continuous Chinese narratives in one ear while simultaneously ignoring a distracting stimulus in the other ear. The results showed equivalent behavioral performance across the two groups, with participants achieving full comprehension across all conditions. However, compared to bilinguals in L1-dominant environment, immersion in an L2 context increased the neural tracking selectivity for attended – but not unattended - stimuli, regardless of the type or presence of interference. This modulation of neural tracking primarily came from the top-down mechanisms captured by the delta frequency band (1-4 Hz), implying that immersion experience modulates higher-order integrative processes and mechanisms associated with difficult listening conditions. Critically, such boost in attentional tracking appeared to decrease following longer L2 immersion, indicative of the flexible tuning of neurocognitive mechanisms as the language environment becomes more stable. Power analyses revealed no group differences in the overall attentional engagement with the listening task, which also confirmed that there were no global spectral and site-related differences between the two immersion groups.

### 4.1 Language immersion shapes how bilinguals attend to speech

Behaviorally, no differences between the immersed and non-immersed group were found in their comprehension performance, and the existence or type of interference did not impact behavioral performance. The high response accuracy showed that both groups understood the attended stories equally well across various listening scenarios, confirming that – unsurprisingly - shifting to an L2-dominant environment did not alter the bilinguals’ overall ability to extract the semantic content from continuous narratives, even in the presence of distractors. On the other hand, the cortical tracking data showed that, despite the striking similarity of the patterns of results across the two groups (Figure 2a), L2 immersion had a clear impact on cortical tracking of attended speech. Here, the immersed bilinguals showed significantly more robust neural tracking of attended speech than their non-immersed counterparts. The tracking of the unattended streams was however near-identical between the two groups. Another notable result from the cortical tracking analyses was that this more robust tracking of the attended streams in the immersed group held across all listening conditions in our experiment, including all interreference conditions, as well as the Single Talker condition.

Previous speech-tracking studies reported that bilingual experience (i.e., L2 exposure, L2 immersion, similarity between L1 and L2) can modulate the neural encoding of attended speech without eliciting behavioral differences between monolinguals and bilinguals (Olguin et al., 2019; Phelps et al., 2023). This was interpreted as indicating that this modulation is adaptive in nature, aimed at enabling optimal behavioral performance in the more linguistically complex processing contexts. More generally, selective attention has been shown to enhance the neural tracking of target speech relative to the unattended input in multi-talker paradigms, with stronger tracking of the attended envelopes reliably associated with greater speech-in-noise understanding (Decruy et al., 2019; Ding & Simon, 2012; Horton et al., 2013; Kurthen et al., 2021; Vanthornhout et al., 2018). Therefore, our finding that L2 immersion boosts cortical tracking of the attended stream suggests that, instead of globally increasing responsiveness to all auditory inputs, immersion in an L2 environment selectively amplifies top-down gain for task-relevant speech signals, arguably reflecting an enhanced “alertness” to target stimuli or information in immersed bilinguals. The result that such effect is most salient in the delta band (1-4 Hz) adds further support to this interpretation, given the widely-reported association between delta-range tracking and top-down processes including syntactic and semantic integration in adverse listening conditions (Etard & Reichenbach, 2019; Ershaid et al., 2024; Ding et al., 2016; Lu et al., 2023; Mai & Wang, 2019; Molinaro & Lizarazu, 2018). The finding that this enhancement applies even to the speech processing in the Single Talker condition, where no overt interference was presented to participants, underlines the powerful impact of language immersion on how bilingual listeners attend to speech signals. This impact would arguably arise due to the constant, high-demand language use that would require understanding and producing L2 in real time and across multiple contexts (social, academic, emotional). Such explicit, effortful processing would increase the need for language control, loading on selection and inhibition mechanisms and the corresponding executive control networks (Alain et al., 2018; Erb & Obleser, 2013). As has been shown with language processing in other challenging linguistic environments (Klimovich-Gray et al., 2026), such contexts can lead to greater reliance on top-down lexico-semantic processing, which is primarily indexed by the delta-range cortical tracking – directly in line with the results observed in the current study.

Finally, it is important to emphasize that these effects emerged while the listeners attended to narratives in their native language, not in L2 as could be more commonly expected (the current study did not test how immersion impacts the neural encoding of the second language). This striking outcome further underlines the impact that immersion has on the way listeners encode speech, and adds strong support to the findings that this reshaping extends beyond the bilinguals’ second language alone (Linck et al., 2009). Taken together, these results indicate that L2 immersion strengthens bilinguals’ capacity to build a robust neural representation of target speech, applying across different listening conditions and likely generalizing across both bilinguals’ languages.

### 4.2 The significance of L2 immersion length

Within the Cambridge group, we observed a significant interaction between L2 immersion time and attention: despite uniformly high comprehension performance, bilinguals with shorter immersion showed larger difference between tracking of attended and unattended streams, with this attentional boost diminishing with longer immersion (although relatively long immersion may be necessary for the effect to emerge, as evidenced by its attenuation following the exclusion of the participant with the longest immersion). This pattern is compatible with accounts that treat bilingual language control as a dynamically adapting system responsive to relative cognitive demands. There have been multiple proposals that language-control processes adapt to the recurrent demands of the interactional context, such that the degree and type of control recruited depend on how taxing the environment is (Bialystok et al., 2009; Green & Abutalebi, 2013; Kroll & Bialystok, 2013). Shift from L1 to L2-speaking country and initial residence in an L2-dominant setting likely impose particularly high demands, with arguably parallel activation of L1 and L2 and strong cross-language competition. The more contemporary neuroplasticity models also predict that such adaptations are not monotonic. DeLuca et al. (2020a, 2020b) conceptualized bilingualism as a trajectory of experience in which factors such as intensity and duration of immersion elicit phase-like modulations in brain structure and function, including periods of expansion and subsequent consolidation. The Dynamic Restructuring Model (DRM) (Pliatsikas, 2020) suggests that experience-based structural adaptations are most pronounced when control demands are high at the early stage and then attenuate as the system reorganizes into a more efficient state, demonstrating non-linear, expansion-renormalization trajectories (e.g., Korenar et al., 2023; Marin-Marín et al., 2022; Wenger et al., 2017b). Our data provide a functional level analogue of such trajectory. In the early and middle stages of L2 immersion, communication in an L2-dominant environment is still cognitively demanding, which requires continuous management of cross-language competition and frequent inhibition of L1 to use L2 (Linck et al., 2009). Thus, when facing challenging listening scenarios requiring sustained attention, the executive function in bilinguals may rely on a strong regulation of top-down gain on the target speech, yielding the pronounced enhancement of attended decoding we observed in immersed bilinguals with fewer years of immersion. As immersion length accumulates and the language environment stabilizes, bilingual language control is expected to become more automated and mechanically efficient (DeLuca et al., 2020a, 2020b; Pliatsikas et al., 2021), and the neural system may no longer need to sustain such elevated levels of cognitive investment in the task-relevant stimuli. From this perspective, the negative association between L2 immersion duration and attended tracking may not mean ability degradation, but rather a renormalization of attentional system once the demands of L2-dominant context become routine and stabilized. Therefore, our findings may extend DRM and the expansion-renormalization framework from structural to task-relevant functional measures. Specifically, the negative association between L2 immersion length and attended tracking supports a dynamic, experience-dependent view of bilingual adaptation, where L2 immersion initially sharpens neural selective attention to speech, but such boosting gradually relaxes as bilinguals move from the early, effortful adaptation stage to a more stable, efficient mode of functioning in the L2-speaking environment.

### 4.3 Immersion does not influence task engagement

The spectral power analyses provide an important complement to the mTRF decoding results. Across the broad 1-12 Hz range and within each decomposed band (delta, theta, alpha), no group or condition differences emerged in the normalized PSD, suggesting that immersed and non-immersed bilingual groups engaged comparable levels of task-related neural resources throughout the entire dichotic listening task. Given that low-frequency power generally reflects large-scale brain engagement of integrative cognitive control, attention and inhibitory processes (e.g., Buzsáki & Draguhn, 2004; Foxe & Snyder, 2011; Pfurtscheller & Lopes da Silva, 1999), the absence of power differences suggests no global group discrepancy in the indices of cognitive control, selective inhibition, attention allocation or listening effort during the task. This pattern of power results complements the observed group differences in mTRF speech tracking, which are confined to attended stimuli in the delta band, collectively suggesting that L2 immersion specifically changes the precision of speech-brain mapping and top-down selective encoding, rather than generic arousal and task-related neural engagement. That is, while both bilingual groups seem to recruit similar amounts of low-frequency neural resources to perform the comprehension listening task, L2 immersion modulates how those resources are functionally deployed, specifically sharpening delta-band synchronization to the target speech stimulus without changing the overall power in the underlying oscillatory systems.

### 4.4 Limitations and open questions

While our results clearly indicate that language immersion modifies attentional processing of speech, with immersion length playing an important role in this process, consistent with the dynamic restructuring model of experience-based skills (e.g., Pliatsikas, 2020; Wenger et al., 2017b), the discontinuous spread of immersion length and the cross-sectional nature of the study only offers snapshots of this trajectory. A critical next step will therefore be longitudinal research that tracks bilinguals across different phases of L2 immersion and a wider spread of immersion length, ideally from pre-departure to several years into residency in the L2-speaking country. Such data would provide a more direct insight into the way neural speech tracking changes over time and whether the attentional gain triggered by L2 immersion follows a systematic expansion-renormalization pattern. This longitudinal design should also incorporate age and L2 AoA as additional continuous predictors, given evidence that AoA may interact with executive outcomes and confound or modulate the influence of immersion (Xia et al., 2025).

In addition, although our participant groups were carefully matched on L2 AoA and proficiency, the immersion indices were necessarily coarse, relying on self-reported duration of residency in the L2-dominant country. In particular, the study did not take into account interactional context (single-language vs dual-language settings) or language entropy, measures that capture the diversity and unpredictability of language use in a particular environment that were shown to be shape control demands and neural adaptation in bilingualism (Gullifer & Titone, 2020). Finally, our study specifically focused on a single language pair (Chinese-English) and on L1 processing alone. Extending this work to other language combinations and systematically comparing attentional processing for L1 and L2 within the same listeners may be essential for determining whether the observed immersion-related modulation is comparable across the bilinguals’ languages. It would also be informative to assess how immersion effects interact with other facets of language experience such as typological distance between L1 and L2, language switching or relative proficiency.

### 4.5 Conclusion

In summary, our research shows that real-world L2 immersion systematically modulates attentional processing of speech in early Mandarin-English bilinguals. Relative to bilinguals living in an L1-dominant environment, those residing in an L2-dominant context exhibited more robust neural tracking of attended speech, driven by the delta band, and consistent across different listening conditions. In addition, we found that prolonged immersion was associated with weaker tracking of the attended signal, suggesting dynamic neurocognitive adaptation of executive mechanisms as the language environment stabilizes. Our data extends ideas of the expansion-renormalization model from structure and resting connectivity to task-dependent, online neural speech tracking. Together with evidence that bilingualism is best understood as a spectrum of experiences leading to a range of neural adaptations (DeLuca et al., 2019a), our study underscores that bilingual modulation of attentional speech processing is not a monotonous, single “bilingual effect”, but a complex multi-dimensional, highly flexible and plastic, dynamically tuned system which is subject to variations of demands exerted by language contexts. Despite no behavioral differences in L1 comprehension, the neural representation of speech under interference is continuously reshaped by experiences such as L2 immersion and its duration even in proficient adult bilinguals. Prolonged L2 exposure was shown to sharpen the brain’s general ability to pick out and follow target auditory information from background noise, with this enhancement appearing to renormalize as immersion duration increases.

## Supporting information

Appendix all tables and figures

## Ethics and integrity statement

The study was approved by the Institutional Review Boards from both Cambridge University and Beijing Normal University.

## Data availability

The datasets and code for this study are available on OSF: https://osf.io/gsrje/

## Acknowledgements

This work was supported by funds from the Department of Psychology, University of Cambridge to JW, Cambridge Language Sciences Incubator award to MB, and National Natural Science Foundation of China (31871097) to TG. We would like to thank Peng Peng and Yanzhuo Li for data collection, and the participants for making this research possible.

## References

Alain, C., Du, Y., Bernstein, L. J., Barten, T., & Banai, K. (2018). Listening under difficult conditions: An activation likelihood estimation meta-analysis. Human brain mapping, 39(7), 2695–2709.

Attaheri, A., Choisdealbha, Á. N., Di Liberto, G. M., Rocha, S., Brusini, P., Mead, N., … & Goswami, U. (2022). Delta-and theta-band cortical tracking and phase-amplitude coupling to sung speech by infants. NeuroImage, 247, 118698.

Bates, D., Mächler, M., Bolker, B., & Walker, S. (2015). Fitting linear mixed-effects models using lme4. Journal of statistical software, 67, 1–48.

Berens, M. S., Kovelman, I., & Petitto, L. A. (2013). Should bilingual children learn reading in two languages at the same time or in sequence?. Bilingual research journal, 36(1), 35–60.

Bialystok, E., Craik, F. I., Green, D. W., & Gollan, T. H. (2009). Bilingual minds. Psychological science in the public interest, 10(3), 89–129.

Birdsong, D., Gertken, L. M., & Amengual, M. (2012). Bilingual language profile: An easy-to-use instrument to assess bilingualism. COERLL, University of Texas at Austin.

Bozic, M., Tyler, L. K., Ives, D. T., Randall, B., & Marslen-Wilson, W. D. (2010). Bihemispheric foundations for human speech comprehension. Proceedings of the National Academy of Sciences, 107(40), 17439–17444.

Brainard, D. H., & Vision, S. (1997). The psychophysics toolbox. Spatial vision, 10(4), 433–436.

Bröhl, F., & Kayser, C. (2021). Delta/theta band EEG differentially tracks low and high frequency speech-derived envelopes. Neuroimage, 233, 117958.

Buzsaki, G., & Draguhn, A. (2004). Neuronal oscillations in cortical networks. science, 304(5679), 1926–1929.

Cavanagh, J. F., & Frank, M. J. (2014). Frontal theta as a mechanism for cognitive control. Trends in cognitive sciences, 18(8), 414–421.

Crosse, M. J., Di Liberto, G. M., Bednar, A., & Lalor, E. C. (2016). The multivariate temporal response function (mTRF) toolbox: a MATLAB toolbox for relating neural signals to continuous stimuli. Frontiers in human neuroscience, 10, 604.

De Cheveigné, A., & Arzounian, D. (2018). Robust detrending, rereferencing, outlier detection, and inpainting for multichannel data. NeuroImage, 172, 903–912.

Decruy, L., Vanthornhout, J., & Francart, T. (2019). Evidence for enhanced neural tracking of the speech envelope underlying age-related speech-in-noise difficulties. Journal of neurophysiology, 122(2), 601–615.

Delorme, A., & Makeig, S. (2004). EEGLAB: an open source toolbox for analysis of single-trial EEG dynamics including independent component analysis. Journal of neuroscience methods, 134(1), 9–21.

DeLuca, V., Rothman, J., & Pliatsikas, C. (2019b). Linguistic immersion and structural effects on the bilingual brain: a longitudinal study. Bilingualism: Language and Cognition, 22(5), 1160–1175.

DeLuca, V., Rothman, J., Bialystok, E., & Pliatsikas, C. (2019a). Redefining bilingualism as a spectrum of experiences that differentially affects brain structure and function. Proceedings of the National Academy of Sciences, 116(15), 7565–7574.

DeLuca, V., Rothman, J., Bialystok, E., & Pliatsikas, C. (2020b). Duration and extent of bilingual experience modulate neurocognitive outcomes. NeuroImage, 204, 116222.

DeLuca, V., Segaert, K., Mazaheri, A., & Krott, A. (2020a). Understanding bilingual brain function and structure changes? U bet! A unified bilingual experience trajectory model. Journal of Neurolinguistics, 56, 100930.

DeLuca, V., Voits, T., Ni, J., Carter, F., Rahman, F., Mazaheri, A., … & Segaert, K. (2024). Mapping individual aspects of bilingual experience to adaptations in brain structure. Cerebral Cortex, 34(2), bhae029.

Dimitrijevic, A., Smith, M. L., Kadis, D. S., & Moore, D. R. (2017). Cortical alpha oscillations predict speech intelligibility. Frontiers in human neuroscience, 11, 88.

Ding, N., & Simon, J. Z. (2012). Neural coding of continuous speech in auditory cortex during monaural and dichotic listening. Journal of neurophysiology, 107(1), 78–89.

Ding, N., Melloni, L., Zhang, H., Tian, X., & Poeppel, D. (2016). Cortical tracking of hierarchical linguistic structures in connected speech. Nature neuroscience, 19(1), 158–164.

Doelling, K. B., Arnal, L. H., Ghitza, O., & Poeppel, D. (2014). Acoustic landmarks drive delta–theta oscillations to enable speech comprehension by facilitating perceptual parsing. Neuroimage, 85, 761–768.

Erb, J., & Obleser, J. (2013). Upregulation of cognitive control networks in older adults’ speech comprehension. Frontiers in systems neuroscience, 7, 116.

Ershaid, H., Lizarazu, M., McLaughlin, D., Cooke, M., Simantiraki, O., Koutsogiannaki, M., & Lallier, M. (2024). Contributions of listening effort and intelligibility to cortical tracking of speech in adverse listening conditions. Cortex, 172, 54–71.

Etard, O., & Reichenbach, T. (2019). Neural speech tracking in the theta and in the delta frequency band differentially encode clarity and comprehension of speech in noise. Journal of Neuroscience, 39(29), 5750–5759.

Foxe, J. J., & Snyder, A. C. (2011). The role of alpha-band brain oscillations as a sensory suppression mechanism during selective attention. Frontiers in psychology, 2, 154.

Frey, J. N., Ruhnau, P., & Weisz, N. (2015). Not so different after all: The same oscillatory processes support different types of attention. Brain research, 1626, 183–197.

Genesee, F. (1987). Learning through two languages: Studies of immersion and bilingual education. (No Title).

Giraud, A. L., & Poeppel, D. (2012). Cortical oscillations and speech processing: emerging computational principles and operations. Nature neuroscience, 15(4), 511–517.

Green, D. W., & Abutalebi, J. (2013). Language control in bilinguals: The adaptive control hypothesis. Journal of cognitive psychology, 25(5), 515–530.

Grundy, J. G., & Timmer, K. (2017). Bilingualism and working memory capacity: A comprehensive meta-analysis. Second Language Research, 33(3), 325–340.

Gullifer, J. W., & Titone, D. (2020). Characterizing the social diversity of bilingualism using language entropy. Bilingualism: Language and cognition, 23(2), 283–294.

Harmony, T. (2013). The functional significance of delta oscillations in cognitive processing. Frontiers in integrative neuroscience, 7, 83.

Horton, C., D’Zmura, M., & Srinivasan, R. (2013). Suppression of competing speech through entrainment of cortical oscillations. Journal of neurophysiology, 109(12), 3082–3093.

Jensen, O., & Mazaheri, A. (2010). Shaping functional architecture by oscillatory alpha activity: gating by inhibition. Frontiers in human neuroscience, 4, 186.

Johnson, R. K., & Swain, M. (Eds.). (1997). Immersion education. Cambridge University Press.

Klimesch, W., Sauseng, P., & Hanslmayr, S. (2007). EEG alpha oscillations: the inhibition–timing hypothesis. Brain research reviews, 53(1), 63–88.

Klimovich-Gray, A., Bozic, M., Molinaro, N., & Lallier, M. (2026). Dyslexia: a window into the cortical mechanisms of adaptive speech analysis. Trends in Neurosciences.

Knyazev, G. G. (2007). Motivation, emotion, and their inhibitory control mirrored in brain oscillations. Neuroscience & Biobehavioral Reviews, 31(3), 377–395.

Korenar, M., Treffers-Daller, J., & Pliatsikas, C. (2023). Dynamic effects of bilingualism on brain structure map onto general principles of experience-based neuroplasticity. Scientific reports, 13(1), 3428.

Kousaie, S., & Phillips, N. A. (2017). A behavioural and electrophysiological investigation of the effect of bilingualism on aging and cognitive control. Neuropsychologia, 94, 23–35.

Kroll, J. F., & Bialystok, E. (2013). Understanding the consequences of bilingualism for language processing and cognition. Journal of cognitive psychology, 25(5), 497–514.

Ktonas, P. Y., & Papp, N. (1980). Instantaneous envelope and phase extraction from real signals: theory, implementation, and an application to EEG analysis. Signal Processing, 2(4), 373–385.

Kurthen, I., Galbier, J., Jagoda, L., Neuschwander, P., Giroud, N., & Meyer, M. (2021). Selective attention modulates neural envelope tracking of informationally masked speech in healthy older adults. Human brain mapping, 42(10), 3042–3057.

Lavie, N. (2010). Attention, distraction, and cognitive control under load. Current directions in psychological science, 19(3), 143–148.

Lemhöfer, K., & Broersma, M. (2012). Introducing LexTALE: A quick and valid lexical test for advanced learners of English. Behavior research methods, 44(2), 325–343.

Leske, S., & Dalal, S. S. (2019). Reducing power line noise in EEG and MEG data via spectrum interpolation. Neuroimage, 189, 763–776.

Linck, J. A., Kroll, J. F., & Sunderman, G. (2009). Losing access to the native language while immersed in a second language: Evidence for the role of inhibition in second-language learning. Psychological science, 20(12), 1507–1515.

Lu, Y., Jin, P., Ding, N., & Tian, X. (2023). Delta-band neural tracking primarily reflects rule-based chunking instead of semantic relatedness between words. Cerebral Cortex, 33(8), 4448–4458.

Luk, G., De Sa, E. R. I. C., & Bialystok, E. (2011). Is there a relation between onset age of bilingualism and enhancement of cognitive control?. Bilingualism: Language and cognition, 14(4), 588–595.

Mai, G., & Wang, W. S. (2019). Delta and theta neural entrainment during phonological and semantic processing in speech perception. BioRxiv, 556837.

Marian, V., Spivey, M., & Hirsch, J. (2003). Shared and separate systems in bilingual language processing: Converging evidence from eyetracking and brain imaging. Brain and language, 86(1), 70–82.

Marin-Marin L, Costumero V, Ávila C and Pliatsikas C (2022). Dynamic Effects of Immersive Bilingualism on Cortical and Subcortical Grey Matter Volumes. Frontiers in Psychology, 13:886222.

Martin, C. D., Dering, B., Thomas, E. M., & Thierry, G. (2009). Brain potentials reveal semantic priming in both the ‘active’and the ‘non-attended’language of early bilinguals. NeuroImage, 47(1), 326–333.

McHaney, J. R., Gnanateja, G. N., Smayda, K. E., Zinszer, B. D., & Chandrasekaran, B. (2021). Cortical tracking of speech in delta band relates to individual differences in speech in noise comprehension in older adults. Ear and Hearing, 42(2), 343–354.

Molinaro, N., & Lizarazu, M. (2018). Delta (but not theta)-band cortical entrainment involves speech-specific processing. European Journal of Neuroscience, 48(7), 2642–2650.

Morales, J., Yudes, C., Gómez-Ariza, C. J., & Bajo, M. T. (2015). Bilingualism modulates dual mechanisms of cognitive control: Evidence from ERPs. Neuropsychologia, 66, 157–169.

Muda, L., Begam, M., & Elamvazuthi, I. (2010). Voice recognition algorithms using mel frequency cepstral coefficient (MFCC) and dynamic time warping (DTW) techniques. arXiv preprint arXiv:1003.4083.

Muthukumaraswamy, S. D. (2013). High-frequency brain activity and muscle artifacts in MEG/EEG: a review and recommendations. Frontiers in human neuroscience, 7, 138.

Nicolay, A.-C., & Poncelet, M. (2013). Cognitive advantage in children enrolled in a second-language immersion elementary school program for three years. Bilingualism: Language and Cognition, 16(3), 597–607.

Obleser, J., & Kayser, C. (2019). Neural entrainment and attentional selection in the listening brain. Trends in cognitive sciences, 23(11), 913–926.

Obleser, J., & Weisz, N. (2012). Suppressed alpha oscillations predict intelligibility of speech and its acoustic details. Cerebral cortex, 22(11), 2466–2477.

Olguin, A., Bekinschtein, T. A., & Bozic, M. (2018). Neural encoding of attended continuous speech under different types of interference. Journal of Cognitive Neuroscience, 30(11), 1606–1619.

Olguin, A., Cekic, M., Bekinschtein, T. A., Katsos, N., & Bozic, M. (2019). Bilingualism and language similarity modify the neural mechanisms of selective attention. Scientific Reports, 9(1), 8204.

Oostenveld, R., Fries, P., Maris, E., & Schoffelen, J. M. (2011). FieldTrip: Open source software for advanced analysis of MEG, EEG, and invasive electrophysiological data. Computational intelligence and neuroscience, 2011(1), 156869.

Pelli, D. G., & Vision, S. (1997). The VideoToolbox software for visual psychophysics: Transforming numbers into movies. Spatial vision, 10, 437–442.

Pfurtscheller, G., & Da Silva, F. L. (1999). Event-related EEG/MEG synchronization and desynchronization: basic principles. Clinical neurophysiology, 110(11), 1842–1857.

Phelps, J. (2023). Neural and behavioural effects of bilingualism on selective attention (Doctoral dissertation).

Phelps, J., & Bozic, M. (2025). Flexible functional adaptation of selective attention in bilingualism. Bilingualism: Language and Cognition, 28(2), 312–326.

Phelps, J., Attaheri, A., & Bozic, M. (2022). How bilingualism modulates selective attention in children. Scientific Reports, 12(1), 6381.

Pliatsikas, C. (2020). Understanding structural plasticity in the bilingual brain: The Dynamic Restructuring Model. Bilingualism: Language and Cognition, 23(2), 459–471.

Pliatsikas, C., DeLuca, V., Moschopoulou, E., & Saddy, J. D. (2017). Immersive bilingualism reshapes the core of the brain. Brain Structure and Function, 222(4), 1785–1795.

Pliatsikas, C., Pereira Soares, S. M., Voits, T., DeLuca, V., & Rothman, J. (2021). Bilingualism is a long-term cognitively challenging experience that modulates metabolite concentrations in the healthy brain. Scientific Reports, 11(1), 7090.

Purić, D., Vuksanović, J., & Chondrogianni, V. (2017). Cognitive advantages of immersion education after 1 year: Effects of amount of exposure. Journal of Experimental Child Psychology, 159, 296–309.

Quallo, M. M., Price, C. J., Ueno, K., Asamizuya, T., Cheng, K., Lemon, R. N., & Iriki, A. (2009). Gray and white matter changes associated with tool-use learning in macaque monkeys. Proceedings of the National Academy of Sciences, 106(43), 18379–18384.

Ruffieux, J., Mouthon, A., Keller, M., Mouthon, M., Annoni, J. M., & Taube, W. (2018). Balance training reduces brain activity during motor simulation of a challenging balance task in older adults: an fMRI study. Frontiers in behavioral neuroscience, 12, 10.

Ruitenberg, M. F. L., Koppelmans, V., De Dios, Y. E., Gadd, N. E., Wood, S. J., Reuter-Lorenz, P. A., … & Seidler, R. D. (2018). Neural correlates of multi-day learning and savings in sensorimotor adaptation. Scientific reports, 8(1), 14286.

Sampaio-Baptista, C., & Johansen-Berg, H. (2017). White matter plasticity in the adult brain. Neuron, 96(6), 1239–1251

Sauseng, P., Griesmayr, B., Freunberger, R., & Klimesch, W. (2010). Control mechanisms in working memory: a possible function of EEG theta oscillations. Neuroscience & Biobehavioral Reviews, 34(7), 1015–1022.

Schlegel, A. A., Rudelson, J. J., & Tse, P. U. (2012). White matter structure changes as adults learn a second language. Journal of cognitive neuroscience, 24(8), 1664–1670.

Starreveld, P. A., De Groot, A. M., Rossmark, B. M., & Van Hell, J. G. (2014). Parallel language activation during word processing in bilinguals: Evidence from word production in sentence context. Bilingualism: Language and Cognition, 17(2), 258–276.

Strauß, A., Wöstmann, M., & Obleser, J. (2014). Cortical alpha oscillations as a tool for auditory selective inhibition. Frontiers in human neuroscience, 8, 350.

Subasi, A., & Gursoy, M. I. (2010). EEG signal classification using PCA, ICA, LDA and support vector machines. Expert systems with applications, 37(12), 8659–8666.

Sullivan, M. D., Janus, M., Moreno, S., Astheimer, L., & Bialystok, E. (2014). Early stage second-language learning improves executive control: Evidence from ERP. Brain and language, 139, 84–98.

Tao, L., Wang, G., Zhu, M., & Cai, Q. (2021). Bilingualism and domain-general cognitive functions from a neural perspective: A systematic review. Neuroscience & Biobehavioral Reviews, 125, 264–295.

Tiwari, V. (2010). MFCC and its applications in speaker recognition. International journal on emerging technologies, 1(1), 19–22.

Uppenkamp, S., Johnsrude, I. S., Norris, D., Marslen-Wilson, W., & Patterson, R. D. (2006). Locating the initial stages of speech–sound processing in human temporal cortex. Neuroimage, 31(3), 1284–1296.

Vanthornhout, J., Decruy, L., Wouters, J., Simon, J. Z., & Francart, T. (2018). Speech intelligibility predicted from neural entrainment of the speech envelope. Journal of the Association for Research in Otolaryngology, 19(2), 181–191.

Wenger, E., Brozzoli, C., Lindenberger, U., & Lövdén, M. (2017b). Expansion and renormalization of human brain structure during skill acquisition. Trends in cognitive sciences, 21(12), 930–939.

Wenger, E., & Kühn, S. (2020). Neuroplasticity. In Cognitive training: An overview of features and applications (pp. 69–83). Cham: Springer International Publishing.

Wenger, E., Kühn, S., Verrel, J., Mårtensson, J., Bodammer, N. C., Lindenberger, U., & Lövdén, M. (2017a). Repeated structural imaging reveals nonlinear progression of experience-dependent volume changes in human motor cortex. Cerebral Cortex, 27(5), 2911–2925.

Whitham, E. M., Pope, K. J., Fitzgibbon, S. P., Lewis, T., Clark, C. R., Loveless, S., … & Willoughby, J. O. (2007). Scalp electrical recording during paralysis: quantitative evidence that EEG frequencies above 20 Hz are contaminated by EMG. Clinical neurophysiology, 118(8), 1877–1888.

Wöstmann, M., Herrmann, B., Maess, B., & Obleser, J. (2016). Spatiotemporal dynamics of auditory attention synchronize with speech. Proceedings of the National Academy of Sciences, 113(14), 3873–3878.

Wöstmann, M., Maess, B., & Obleser, J. (2021). Orienting auditory attention in time: Lateralized alpha power reflects spatio-temporal filtering. NeuroImage, 228, 117711.

Xia, L., Sorace, A., Vega-Mendoza, M., & Bak, T. (2025). How language proficiency and age of acquisition affect executive control in bilinguals: continuous versus dichotomous analysis approaches. Bilingualism: Language and Cognition, 1–14.

Xie, Z., & Dong, Y. (2021). Influence of the study abroad bilingual experience on cognitive control among young adults. International Journal of Bilingualism, 25(5), 1417–1428.

Yow, W. Q., & Li, X. (2015). Balanced bilingualism and early age of second language acquisition as the underlying mechanisms of a bilingual executive control advantage: Why variations in bilingual experiences matter. Frontiers in psychology, 6, 164.

Zatorre, R. J., Fields, R. D., & Johansen-Berg, H. (2012). Plasticity in gray and white: neuroimaging changes in brain structure during learning. Nature neuroscience, 15(4), 528–536.

Zuk, N. J., Murphy, J. W., Reilly, R. B., & Lalor, E. C. (2021). Envelope reconstruction of speech and music highlights stronger tracking of speech at low frequencies. PLoS computational biology, 17(9), e1009358.

