## Appendix all tables and figures for "Language Immersion Enhances Attentional Speech Processing via Low-Frequency Neural Tracking"

**Table S1**. BLP questionnaire responses for Cambridge and Beijing participants

| **Group** | **Cambridge** | | **Beijing** | |
|---|---|---|---|---|
| ***Language history*** | ***Chinese*** | ***English*** | ***Chinese*** | ***English*** |
| At what age did you start learning the following languages? | M=0.19, SD=0.40 | M=4.89, SD=1.05 | M=0.55, SD=0.89 | M=4.52, SD=1.36 |
| At what age did you start to feel comfortable using the following languages? | M=0.89, SD=1.34 | M=13.00, SD=3.98 | M=2.32, SD=2.12 | M=8.48, SD=4.43 |
| How many years of course have you had in the following languages (primary school through university)? | M=15.30, SD=4.27 | M=11.93, SD=5.22 | M=15.87, SD=3.62 | M= 14.16, SD=3.03 |
| How many years have you spent in a country/region where the following languages are spoken? | M=21.11, SD=3.76 | M=3.22, SD=2.81 | M=22.13, SD=2.75 | M=0.23, SD=0.50 |
| How many years have you spent in a family where the following languages are spoken? | M=22.52, SD=3.15 | M=0.52, SD=1.31 | M=22.16, SD=2.75 | M=0.16, SD=0.52 |
| How many years have you spent in a work environment where the following languages are spoken? | M=7.63, SD=10.74 | M=1.56, SD=1.89 | M=12.23, SD=10.71 | M=1.74, SD=2.98 |
| ***Language proficiency*** | ***Chinese*** | ***English*** | ***Chinese*** | ***English*** |
| How well do you speak Chinese /English? | 6.00 | 4.63 | 5.68 | 4.55 |
| How well do you understand Chinese /English? | 5.96 | 4.81 | 5.65 | 4.55 |
| How well do you read Chinese /English? | 6.00 | 5.15 | 5.65 | 4.68 |
| How well do you write Chinese /English? | 5.89 | 4.89 | 5.26 | 4.42 |
| ***Language Usage*** | ***Chinese*** | ***English*** | ***Chinese*** | ***English*** |
| In an average week, what % of the time do you use the following languages with friends? | 64.07% | 35.93% | 81.29% | 18.71% |
| In an average week, what % of the time do you use the following languages with family? | 94.81% | 4.44% | 96.45% | 3.55% |
| In an average week, what % of the time do you use the following languages at school/work? | 17.78% | 81.85% | 71.61% | 28.06% |
| When you talk to yourself, how often do you talk to yourself in the following languages? | 73.70% | 25.19% | 76.45% | 23.23% |
| When you count, how often do you count in the following languages? | 77.78% | 19.63% | 81.94% | 17.42% |

**Table S2**. Cortical Speech Tracking: Broadband (1-12 Hz)

*r ~ Attention*Group*Condition + (1|Participant)*

| Effect | Sum Sq | Mean Sq | NumDF | DenDF | F value | Pr(>F) | Eta2_partial |
|---|---|---|---|---|---|---|---|
| Attention | 0.178 | 0.178 | 1 | 280 | 430.603 | **<0.001** | 0.606 |
| Group | 0.005 | 0.005 | 1 | 56 | 12.532 | **<0.001** | 0.183 |
| Condition | 0.002 | 0.001 | 2 | 280 | 2.594 | 0.077 | 0.018 |
| Attention:Group | 0.012 | 0.012 | 1 | 280 | 29.472 | **<0.001** | 0.095 |
| Attention:Condition | 0.026 | 0.013 | 2 | 280 | 31.204 | **<0.001** | 0.182 |
| Group:Condition | 0.0007 | 0.0003 | 2 | 280 | 0.807 | 0.447 | 0.006 |
| Attention:Group:Condition | 0.001 | 0.0005 | 2 | 280 | 1.285 | 0.278 | 0.009 |

*Post-Hoc effects (Tukey corrected):*

| **Attention Effect (Attended vs Unattended)** | | | | | | |
|---|---|---|---|---|---|---|
| ***Group*** | ***Condition*** | ***estimate*** | ***SE*** | ***df*** | ***t.ratio*** | ***p.value*** |
| Cambridge | Native-MuR | 0.085 | 0.006 | 280 | 15.301 | **<0.001** |
| | Native-Unknown | 0.035 | 0.006 | 280 | 6.413 | **<0.001** |
| | Native-Native | 0.051 | 0.006 | 280 | 9.298 | **<0.001** |
| Beijing | Native-MuR | 0.051 | 0.005 | 280 | 9.885 | **<0.001** |
| | Native-Unknown | 0.016 | 0.005 | 280 | 3.166 | **0.002** |
| | Native-Native | 0.033 | 0.005 | 280 | 6.398 | **<0.001** |
| **Group Effect (Cambridge vs Beijing)** | | | | | | |
| ***Attention*** | ***Condition*** | ***estimate*** | ***SE*** | ***df*** | ***t.ratio*** | ***p.value*** |
| Attended | Native-MuR | 0.035 | 0.006 | 223.612 | 5.466 | **<0.001** |
| | Native-Unknown | 0.024 | 0.006 | 223.612 | 3.756 | **<0.001** |
| | Native-Native | 0.021 | 0.006 | 223.612 | 3.244 | **0.001** |
| Unattended | Native-MuR | 0.002 | 0.006 | 223.612 | 0.274 | 0.784 |
| | Native-Unknown | 0.005 | 0.006 | 223.612 | 0.801 | 0.424 |
| | Native-Native | 0.003 | 0.006 | 223.612 | 0.401 | 0.689 |

**Table S3**. Cortical Speech Tracking in Attended Streams only (1-12 Hz)

*Attended r ~ Group*Condition + (1|Participant)*

| Effect | Sum Sq | Mean Sq | NumDF | DenDF | F value | Pr(>F) | Eta2_partial |
|---|---|---|---|---|---|---|---|
| Group | 0.007 | 0.007 | 1 | 56 | 17.446 | **<0.001** | 0.238 |
| Condition | 0.034 | 0.011 | 3 | 168 | 26.470 | **<0.001** | 0.321 |
| Group: Condition | 0.002 | 0.0006 | 3 | 168 | 1.298 | 0.277 | 0.023 |

*Post-Hoc effects (Tukey corrected):*

| **Group Effect (Cambridge vs Beijing)** | | | | | |
|---|---|---|---|---|---|
| ***Condition*** | ***estimate*** | ***SE*** | ***df*** | ***t.ratio*** | ***p.value*** |
| Single Talker | 0.028 | 0.008 | 118.692 | 3.534 | **<0.001** |
| Native-MuR | 0.035 | 0.008 | 118.692 | 4.392 | **<0.001** |
| Native-Unknown | 0.024 | 0.008 | 118.692 | 3.018 | **0.003** |
| Native-Native | 0.021 | 0.008 | 118.692 | 2.607 | **0.010** |

| **Condition Effect (Pairwise)** | | | | | | |
|---|---|---|---|---|---|---|
| ***Group*** | ***Condition contrast*** | ***estimate*** | ***SE*** | ***df*** | ***t.ratio*** | ***p.value*** |
| Cambridge | ST vs Native-MuR | 0.001 | 0.006 | 168 | 0.212 | 0.997 |
| | ST vs Native-Unknown | 0.033 | 0.006 | 168 | 5.891 | **<0.001** |
| | ST vs Native-Native | 0.022 | 0.006 | 168 | 3.833 | **0.001** |
| | Nat-MuR vs Nat-Unknow | 0.032 | 0.006 | 168 | 5.680 | **<0.001** |
| | Nat-MuR vs Nat-Nat | 0.020 | 0.006 | 168 | 3.621 | **0.002** |

| | | | | | | |
|---|---|---|---|---|---|---|
| | Nat-Unknow vs Nat-Nat | -0.012 | 0.006 | 168 | -2.059 | 0.171 |
| Beijing | ST vs Native-MuR | 0.008 | 0.005 | 168 | 1.543 | 0.414 |
| | ST vs Native-Unknown | 0.029 | 0.005 | 168 | 5.520 | **<0.001** |
| | ST vs Native-Native | 0.014 | 0.005 | 168 | 2.684 | **0.040** |
| | Nat-MuR vs Nat-Unknow | 0.021 | 0.005 | 168 | 3.977 | **<0.001** |
| | Nat-MuR vs Nat-Nat | 0.006 | 0.005 | 168 | 1.141 | 0.665 |
| | Nat-Unknow vs Nat-Nat | -0.015 | 0.005 | 168 | -2.836 | **0.026** |

**Table S4.** Cortical Speech Tracking in Unattended Streams only (1-12 Hz)

*Unattended r ~ Group*Condition + (1|Participant)*

| Effect | Sum Sq | Mean Sq | NumDF | DenDF | F value | Pr(>F) | Eta2_partial |
|---|---|---|---|---|---|---|---|
| Group | 0.0003 | 0.0003 | 1 | 56 | 0.989 | 0.324 | 0.017 |
| Condition | 0.008 | 0.004 | 2 | 112 | 13.574 | **<0.001** | 0.195 |
| Group:Condition | 0.00009 | 0.00005 | 2 | 112 | 0.159 | 0.853 | 0.003 |

*Post-Hoc effects (Tukey corrected):*

| **Group Effect (Cambridge vs Beijing)** | | | | | |
|---|---|---|---|---|---|
| ***Condition*** | ***estimate*** | ***SE*** | ***df*** | ***t.ratio*** | ***p.value*** |
| Native-MuR | 0.002 | 0.005 | 160.276 | 0.366 | 0.715 |
| Native-Unknown | 0.005 | 0.005 | 160.276 | 1.069 | 0.287 |
| Native-Native | 0.003 | 0.005 | 160.276 | 0.536 | 0.593 |

| **Condition Effect (Pairwise)** | | | | | | |
|---|---|---|---|---|---|---|
| ***Group*** | ***Condition contrast*** | ***estimate*** | ***SE*** | ***df*** | ***t.ratio*** | ***p.value*** |
| Cambridge | Nat-MuR vs Nat-Unknow | -0.017 | 0.005 | 112 | -3.736 | **<0.001** |
| | Nat-MuR vs Nat-Nat | -0.013 | 0.005 | 112 | -2.785 | **0.017** |
| | Nat-Unknow vs Nat-Nat | 0.004 | 0.005 | 112 | 0.950 | 0.610 |
| Beijing | Nat-MuR vs Nat-Unknow | -0.014 | 0.004 | 112 | -3.210 | **0.005** |
| | Nat-MuR vs Nat-Nat | -0.012 | 0.004 | 112 | -2.793 | **0.017** |
| | Nat-Unknow vs Nat-Nat | 0.002 | 0.004 | 112 | 0.417 | 0.909 |

**Table S5**. Cortical Speech Tracking: Delta band (1-4 Hz)

*r ~ Attention*Group*Condition + (1|Participant)*

| Effect | Sum Sq | Mean Sq | NumDF | DenDF | F value | Pr(>F) | Eta2_partial |
|---|---|---|---|---|---|---|---|
| Attention | 0.154 | 0.154 | 1 | 280 | 347.587 | **<0.001** | 0.554 |
| Group | 0.005 | 0.005 | 1 | 56 | 11.221 | **0.001** | 0.167 |
| Condition | 0.0005 | 0.0002 | 2 | 280 | 0.558 | 0.573 | 0.004 |
| Attention:Group | 0.012 | 0.012 | 1 | 280 | 26.981 | **<0.001** | 0.088 |
| Attention:Condition | 0.016 | 0.008 | 2 | 280 | 18.351 | **<0.001** | 0.116 |

| | | | | | | | |
|---|---|---|---|---|---|---|---|
| Group:Condition | 0.001 | 0.0007 | 2 | 280 | 1.636 | 0.197 | 0.012 |
| Attention:Group:Condition | 0.0004 | 0.0002 | 2 | 280 | 0.501 | 0.607 | 0.004 |

*Post-Hoc effects (Tukey corrected):*

| **Attention Effect (Attended vs Unattended)** | | | | | | |
|---|---|---|---|---|---|---|
| ***Group*** | ***Condition*** | ***estimate*** | ***SE*** | ***df*** | ***t.ratio*** | ***p.value*** |
| Cambridge | Native-MuR | 0.075 | 0.006 | 280 | 13.158 | **<0.001** |
| | Native-Unknown | 0.038 | 0.006 | 280 | 6.718 | **<0.001** |
| | Native-Native | 0.048 | 0.006 | 280 | 8.362 | **<0.001** |
| Beijing | Native-MuR | 0.046 | 0.005 | 280 | 8.543 | **<0.001** |
| | Native-Unknown | 0.017 | 0.005 | 280 | 3.128 | **0.002** |
| | Native-Native | 0.029 | 0.005 | 280 | 5.401 | **<0.001** |
| **Group Effect (Cambridge vs Beijing)** | | | | | | |
| ***Attention*** | ***Condition*** | ***estimate*** | ***SE*** | ***df*** | ***t.ratio*** | ***p.value*** |
| Attended | Native-MuR | 0.033 | 0.007 | 240.734 | 5.096 | **<0.001** |
| | Native-Unknown | 0.026 | 0.007 | 240.734 | 3.902 | **<0.001** |
| | Native-Native | 0.018 | 0.007 | 240.734 | 2.759 | **0.006** |
| Unattended | Native-MuR | 0.004 | 0.007 | 240.734 | 0.552 | 0.582 |
| | Native-Unknown | 0.004 | 0.007 | 240.734 | 0.572 | 0.568 |
| | Native-Native | -0.001 | 0.007 | 240.734 | -0.152 | 0.879 |

**Table S6**. Cortical Speech Tracking in Attended Streams only (1-4 Hz)

*Attended r ~ Group*Condition + (1|Participant)*

| Effect | Sum Sq | Mean Sq | NumDF | DenDF | F value | Pr(>F) | Eta2_partial |
|---|---|---|---|---|---|---|---|
| Group | 0.006 | 0.006 | 1 | 56 | 16.147 | **<0.001** | 0.224 |
| Condition | 0.020 | 0.007 | 3 | 168 | 17.401 | **<0.001** | 0.237 |
| Group: Condition | 0.002 | 0.0006 | 3 | 168 | 1.622 | 0.186 | 0.028 |

*Post-Hoc effects (Tukey corrected):*

| **Group Effect (Cambridge vs Beijing) in Attended Streams** | | | | | | |
|---|---|---|---|---|---|---|
| ***Condition*** | | ***estimate*** | ***SE*** | ***df*** | ***t.ratio*** | ***p.value*** |
| Single Talker | | 0.030 | 0.008 | 110.960 | 3.729 | **<0.001** |
| Native-MuR | | 0.033 | 0.008 | 110.960 | 4.158 | **<0.001** |
| Native-Unknown | | 0.026 | 0.008 | 110.960 | 3.184 | **0.002** |
| Native-Native | | 0.018 | 0.008 | 110.960 | 2.252 | **0.026** |
| **Condition Effect (Pairwise)** | | | | | | |
| ***Group*** | ***Condition contrast*** | ***estimate*** | ***SE*** | ***df*** | ***t.ratio*** | ***p.value*** |
| Cambridge | ST vs Native-MuR | 0.005 | 0.005 | 168 | 0.885 | 0.813 |
| | ST vs Native-Unknown | 0.027 | 0.005 | 168 | 4.994 | **<0.001** |
| | ST vs Native-Native | 0.022 | 0.005 | 168 | 4.138 | **<0.001** |

| | | | | | | |
|---|---|---|---|---|---|---|
| | Nat-MuR vs Nat-Unknow | 0.022 | 0.005 | 168 | 4.109 | **<0.001** |
| | Nat-MuR vs Nat-Nat | 0.017 | 0.005 | 168 | 3.252 | **0.007** |
| | Nat-Unknow vs Nat-Nat | -0.005 | 0.005 | 168 | -0.856 | 0.827 |
| Beijing | ST vs Native-MuR | 0.008 | 0.005 | 168 | 1.637 | 0.361 |
| | ST vs Native-Unknown | 0.022 | 0.005 | 168 | 4.476 | **<0.001** |
| | ST vs Native-Native | 0.010 | 0.005 | 168 | 2.063 | 0.170 |
| | Nat-MuR vs Nat-Unknow | 0.014 | 0.005 | 168 | 2.839 | **0.026** |
| | Nat-MuR vs Nat-Nat | 0.002 | 0.005 | 168 | 0.426 | 0.974 |
| | Nat-Unknow vs Nat-Nat | -0.012 | 0.005 | 168 | -2.413 | 0.078 |

**Table S7.** Cortical Speech Tracking in Unattended Streams only (1-4 Hz)

*Unattended r ~ Group*Condition + (1|Participant)*

| Effect | Sum Sq | Mean Sq | NumDF | DenDF | F value | Pr(>F) | Eta2_partial |
|---|---|---|---|---|---|---|---|
| Group | 0.0001 | 0.0001 | 1 | 56 | 0.405 | 0.527 | 0.007 |
| Condition | 0.007 | 0.004 | 2 | 112 | 12.150 | **<0.001** | 0.178 |
| Group:Condition | 0.0002 | 0.0001 | 2 | 112 | 0.348 | 0.707 | 0.006 |

*Post-Hoc effects (Tukey corrected):*

| **Group Effect (Cambridge vs Beijing)** | | | | | |
|---|---|---|---|---|---|
| ***Condition*** | ***estimate*** | ***SE*** | ***df*** | ***t.ratio*** | ***p.value*** |
| Native-MuR | 0.004 | 0.005 | 159.324 | 0.722 | 0.471 |
| Native-Unknown | 0.004 | 0.005 | 159.324 | 0.748 | 0.455 |
| Native-Native | -0.001 | 0.005 | 159.324 | -0.199 | 0.843 |

| **Condition Effect (Pairwise)** | | | | | | |
|---|---|---|---|---|---|---|
| ***Group*** | ***Condition contrast*** | ***estimate*** | ***SE*** | ***df*** | ***t.ratio*** | ***p.value*** |
| Cambridge | Nat-MuR vs Nat-Unknow | -0.015 | 0.005 | 112 | -3.160 | **0.006** |
| | Nat-MuR vs Nat-Nat | -0.010 | 0.005 | 112 | -2.135 | 0.087 |
| | Nat-Unknow vs Nat-Nat | 0.005 | 0.005 | 112 | 1.025 | 0.563 |
| Beijing | Nat-MuR vs Nat-Unknow | -0.015 | 0.004 | 112 | -3.356 | **0.003** |
| | Nat-MuR vs Nat-Nat | -0.015 | 0.004 | 112 | -3.332 | **0.003** |
| | Nat-Unknow vs Nat-Nat | 0.0001 | 0.004 | 112 | 0.024 | >0.999 |

**Table S8**. Cortical Speech Tracking: Theta band (4-8 Hz)

*r ~ Attention*Group*Condition + (1|Participant)*

| Effect | Sum Sq | Mean Sq | NumDF | DenDF | F value | Pr(>F) | Eta2_partial |
|---|---|---|---|---|---|---|---|
| Attention | 0.009 | 0.009 | 1 | 280 | 55.150 | **<0.001** | 0.165 |
| Group | 0.0006 | 0.0006 | 1 | 56 | 3.335 | 0.073 | 0.056 |
| Condition | 0.0007 | 0.0004 | 2 | 280 | 2.092 | 0.125 | 0.015 |

| | | | | | | | |
|---|---|---|---|---|---|---|---|
| Attention:Group | 0.0007 | 0.0007 | 1 | 280 | 4.149 | **0.043** | 0.015 |
| Attention:Condition | 0.003 | 0.002 | 2 | 280 | 8.824 | **<0.001** | 0.059 |
| Group:Condition | 0.00001 | 0.000007 | 2 | 280 | 0.043 | 0.958 | 0.0003 |
| Attention:Group:Condition | 0.001 | 0.0006 | 2 | 280 | 3.390 | **0.035** | 0.024 |

*Post-Hoc effects (Tukey corrected):*

| **Attention Effect (Attended vs Unattended)** | | | | | | |
|---|---|---|---|---|---|---|
| ***Group*** | ***Condition*** | ***estimate*** | ***SE*** | ***df*** | ***t.ratio*** | ***p.value*** |
| Cambridge | Native-MuR | 0.017 | 0.004 | 280 | 4.839 | **<0.001** |
| | Native-Unknown | 0.003 | 0.004 | 280 | 0.837 | 0.403 |
| | Native-Native | 0.020 | 0.004 | 280 | 5.534 | **<0.001** |
| Beijing | Native-MuR | 0.017 | 0.003 | 280 | 5.038 | **<0.001** |
| | Native-Unknown | 0.002 | 0.003 | 280 | 0.713 | 0.477 |
| | Native-Native | 0.004 | 0.003 | 280 | 1.090 | 0.277 |
| **Group Effect (Cambridge vs Beijing)** | | | | | | |
| ***Attention*** | ***Condition*** | ***estimate*** | ***SE*** | ***df*** | ***t.ratio*** | ***p.value*** |
| Attended | Native-MuR | 0.004 | 0.004 | 329.408 | 1.020 | 0.308 |
| | Native-Unknown | 0.003 | 0.004 | 329.408 | 0.778 | 0.437 |
| | Native-Native | 0.011 | 0.004 | 329.408 | 3.183 | **0.002** |
| Unattended | Native-MuR | 0.003 | 0.004 | 329.408 | 0.883 | 0.378 |
| | Native-Unknown | 0.002 | 0.004 | 329.408 | 0.606 | 0.545 |
| | Native-Native | -0.005 | 0.004 | 329.408 | -1.337 | 0.182 |

**Table S9**. Cortical Speech Tracking in Attended Streams only (4-8 Hz)

*Attended r ~ Group*Condition + (1|Participant)*

| Effect | Sum Sq | Mean Sq | NumDF | DenDF | F value | Pr(>F) | Eta2_partial |
|---|---|---|---|---|---|---|---|
| Group | 0.0002 | 0.0002 | 1 | 56 | 0.546 | 0.463 | 0.010 |
| Condition | 0.012 | 0.004 | 3 | 168 | 11.873 | **<0.001** | 0.175 |
| Group: Condition | 0.003 | 0.001 | 3 | 168 | 3.320 | **0.021** | 0.056 |

*Post-Hoc effects (Tukey corrected):*

| **Group Effect (Cambridge vs Beijing) in Attended Streams** | | | | | | |
|---|---|---|---|---|---|---|
| ***Condition*** | | ***estimate*** | ***SE*** | ***df*** | ***t.ratio*** | ***p.value*** |
| Single Talker | | -0.010 | 0.005 | 219.247 | -1.926 | 0.055 |
| Native-MuR | | 0.004 | 0.005 | 219.247 | 0.733 | 0.464 |
| Native-Unknown | | 0.003 | 0.005 | 219.247 | 0.559 | 0.576 |
| Native-Native | | 0.011 | 0.005 | 219.247 | 2.289 | **0.023** |
| **Condition Effect (Pairwise)** | | | | | | |
| ***Group*** | ***Condition contrast*** | ***estimate*** | ***SE*** | ***df*** | ***t.ratio*** | ***p.value*** |
| Cambridge | ST vs Native-MuR | 0.003 | 0.005 | 168 | 0.625 | 0.924 |

| | | | | | | |
|---|---|---|---|---|---|---|
| | ST vs Native-Unknown | 0.014 | 0.005 | 168 | 2.782 | **0.030** |
| | ST vs Native-Native | 0.002 | 0.005 | 168 | 0.351 | 0.985 |
| | Nat-MuR vs Nat-Unknow | 0.011 | 0.005 | 168 | 2.157 | 0.140 |
| | Nat-MuR vs Nat-Nat | -0.001 | 0.005 | 168 | -0.274 | 0.993 |
| | Nat-Unknow vs Nat-Nat | -0.012 | 0.005 | 168 | -2.431 | 0.075 |
| Beijing | ST vs Native-MuR | 0.016 | 0.005 | 168 | 3.551 | **0.003** |
| | ST vs Native-Unknown | 0.026 | 0.005 | 168 | 5.674 | **<0.001** |
| | ST vs Native-Native | 0.023 | 0.005 | 168 | 4.943 | **<0.001** |
| | Nat-MuR vs Nat-Unknow | 0.010 | 0.005 | 168 | 2.123 | 0.150 |
| | Nat-MuR vs Nat-Nat | 0.006 | 0.005 | 168 | 1.392 | 0.506 |
| | Nat-Unknow vs Nat-Nat | -0.003 | 0.005 | 168 | -0.731 | 0.884 |

**Table S10.** Cortical Speech Tracking in Unattended Streams only (4-8 Hz)

*Unattended r ~ Group*Condition + (1|Participant)*

| Effect | Sum Sq | Mean Sq | NumDF | DenDF | F value | Pr(>F) | Eta2_partial |
|---|---|---|---|---|---|---|---|
| Group | 0.000001 | 0.000001 | 1 | 168 | 0.009 | 0.925 | 0.00005 |
| Condition | 0.0005 | 0.0003 | 2 | 168 | 1.681 | 0.189 | 0.020 |
| Group:Condition | 0.0005 | 0.0003 | 2 | 168 | 1.711 | 0.184 | 0.020 |

*Post-Hoc effects (Tukey corrected):*

| **Group Effect (Cambridge vs Beijing)** | | | | | |
|---|---|---|---|---|---|
| ***Condition*** | ***estimate*** | ***SE*** | ***df*** | ***t.ratio*** | ***p.value*** |
| Native-MuR | 0.003 | 0.003 | 168 | 0.955 | 0.341 |
| Native-Unknown | 0.002 | 0.003 | 168 | 0.655 | 0.513 |
| Native-Native | -0.005 | 0.003 | 168 | -1.446 | 0.150 |

| **Condition Effect (Pairwise)** | | | | | | |
|---|---|---|---|---|---|---|
| ***Group*** | ***Condition contrast*** | ***estimate*** | ***SE*** | ***df*** | ***t.ratio*** | ***p.value*** |
| Cambridge | Nat-MuR vs Nat-Unknow | -0.004 | 0.003 | 112 | -1.085 | 0.525 |
| | Nat-MuR vs Nat-Nat | 0.001 | 0.003 | 112 | 0.334 | 0.940 |
| | Nat-Unknow vs Nat-Nat | 0.005 | 0.003 | 112 | 1.419 | 0.335 |
| Beijing | Nat-MuR vs Nat-Unknow | -0.005 | 0.003 | 112 | -1.473 | 0.308 |
| | Nat-MuR vs Nat-Nat | -0.007 | 0.003 | 112 | -2.130 | 0.088 |
| | Nat-Unknow vs Nat-Nat | -0.002 | 0.003 | 112 | -0.657 | 0.789 |

**Table S11.** Effects of L2 Immersion Length: Broadband (1-12 Hz), Immersed group

*r ~ Immersion time*Condition* Attention + (1|Participant)*

| Effect | Sum Sq | Mean Sq | NumDF | DenDF | F value | Pr(>F) | Eta2_partial |
|---|---|---|---|---|---|---|---|
| Condition | 0.001 | 0.0006 | 2 | 125 | 1.300 | 0.276 | 0.020 |
| Attention | 0.077 | 0.077 | 1 | 125 | 171.166 | **< 0.001** | 0.578 |
| Immersion time | 0.0005 | 0.0005 | 1 | 25 | 1.077 | 0.309 | 0.041 |
| Condition: Attention | 0.007 | 0.004 | 2 | 125 | 8.047 | **<0.001** | 0.114 |
| Condition: Immersion time | 0.0004 | 0.0002 | 2 | 125 | 0.420 | 0.658 | 0.007 |
| Attention: Immersion time | 0.003 | 0.003 | 1 | 125 | 6.249 | **0.014** | 0.048 |
| Condition: Attention: Immersion time | 0.00008 | 0.00004 | 2 | 125 | 0.090 | 0.914 | 0.001 |

**Table S12.** Effects of L2 Immersion Length: Broadband (1-12 Hz), Immersed group, Participant with longest immersion removed

*r ~ Immersion time*Condition* Attention + (1|Participant)*

| Effect | Sum Sq | Mean Sq | NumDF | DenDF | F value | Pr(>F) | Eta2_partial |
|---|---|---|---|---|---|---|---|
| Condition | 0.001 | 0.0005 | 2 | 120 | 1.210 | 0.302 | 0.020 |
| Attention | 0.043 | 0.043 | 1 | 120 | 95.207 | **<0.001** | 0.442 |
| Immersion time | 0.0003 | 0.0003 | 1 | 24 | 0.678 | 0.418 | 0.027 |
| Condition: Attention | 0.006 | 0.003 | 2 | 120 | 6.299 | **0.003** | 0.095 |
| Condition: Immersion time | 0.0007 | 0.0004 | 2 | 120 | 0.814 | 0.446 | 0.013 |
| Attention: Immersion time | 0.0001 | 0.0001 | 1 | 120 | 0.230 | 0.633 | 0.002 |
| Condition: Attention: Immersion time | 0.0003 | 0.0002 | 2 | 120 | 0.335 | 0.716 | 0.006 |

**Figure S1.** The relationship between r Speech Tracking and L2 Immersion time, participant with longest immersion removed

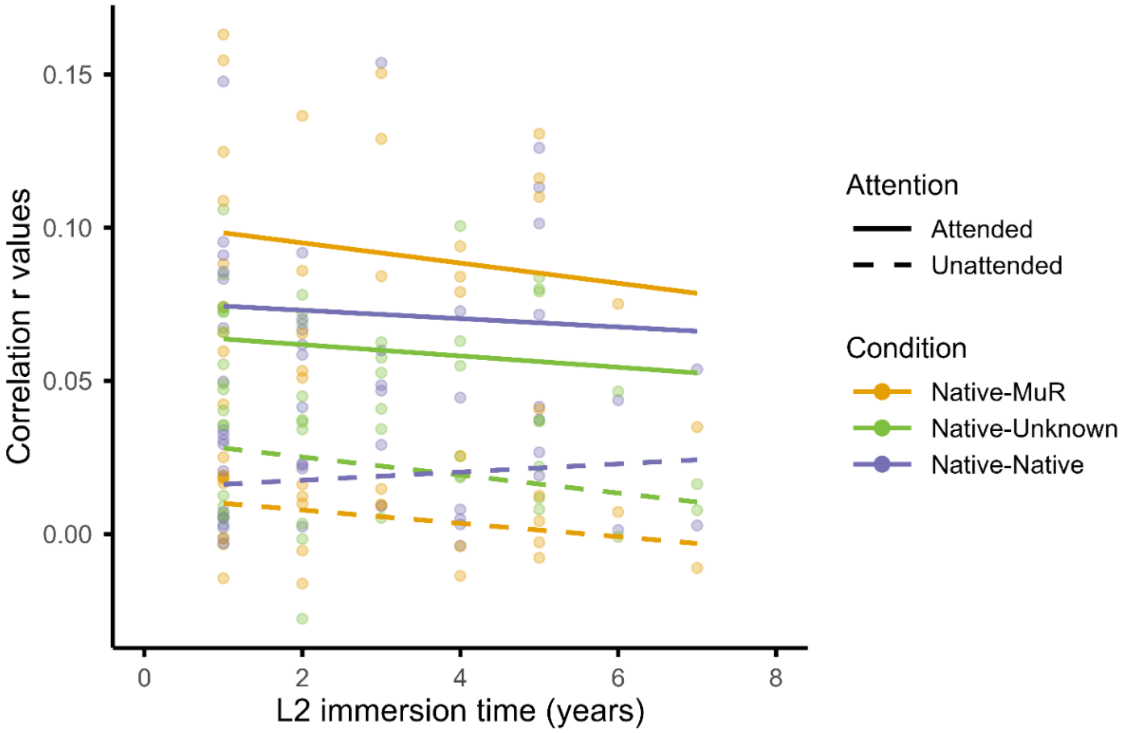


**Table S13.** Power Spectral Density Analysis

*EEG Power ~ Group*Condition + (1|Participant)*

| **Broadband (1-12Hz)** | | | | | | | |
|---|---|---|---|---|---|---|---|
| Effect | Sum Sq | Mean Sq | NumDF | DenDF | F value | Pr(>F) | Eta2_partial |
| Group | 1.166 | 1.166 | 1 | 56 | 0.392 | 0.534 | 0.007 |
| Condition | 7.673 | 2.558 | 3 | 632 | 0.861 | 0.461 | 0.004 |
| Group: Condition | 14.351 | 4.784 | 3 | 632 | 1.610 | 0.186 | 0.008 |
| **Delta band (1-4Hz)** | | | | | | | |
| Effect | Sum Sq | Mean Sq | NumDF | DenDF | F value | Pr(>F) | Eta2_partial |
| Group | 0.012 | 0.012 | 1 | 56 | 0.002 | 0.961 | 0.00004 |
| Condition | 11.036 | 3.679 | 3 | 168 | 0.745 | 0.527 | 0.013 |
| Group: Condition | 20.553 | 6.851 | 3 | 168 | 1.387 | 0.249 | 0.024 |
| **Theta band (4-8Hz)** | | | | | | | |
| Effect | Sum Sq | Mean Sq | NumDF | DenDF | F value | Pr(>F) | Eta2_partial |
| Group | 0.034 | 0.034 | 1 | 56 | 0.405 | 0.527 | 0.007 |
| Condition | 0.642 | 0.214 | 3 | 168 | 2.567 | 0.056 | 0.044 |
| Group: Condition | 0.516 | 0.172 | 3 | 168 | 2.061 | 0.107 | 0.035 |
| **Alpha band (8-12Hz)** | | | | | | | |
| Effect | Sum Sq | Mean Sq | NumDF | DenDF | F value | Pr(>F) | Eta2_partial |
| Group | 1.474 | 1.474 | 1 | 56 | 0.842 | 0.363 | 0.015 |
| Condition | 2.541 | 0.847 | 3 | 168 | 0.484 | 0.694 | 0.009 |
| Group: Condition | 4.483 | 1.494 | 3 | 168 | 0.853 | 0.467 | 0.015 |
